# Non-CG methylation and multiple epigenetic layers associate child abuse with immune and small GTPase dysregulation

**DOI:** 10.1101/501239

**Authors:** Pierre-Eric Lutz, Marc-Aurèle Chay, Alain Pacis, Gary G Chen, Zahia Aouabed, Elisabetta Maffioletti, Jean-François Théroux, Jean-Christophe Grenier, Jennie Yang, Maria Aguirre, Carl Ernst, Adriana Redensek, Léon C. van Kempen, Ipek Yalcin, Tony Kwan, Naguib Mechawar, Tomi Pastinen, Gustavo Turecki

## Abstract

Early-life adversity (ELA) is a major predictor of psychopathology, and is thought to increase lifetime risk by epigenetically regulating the genome. Here, focusing on the lateral amygdala, a major brain site for emotional homeostasis, we described molecular cross-talk among multiple epigenetic mechanisms, including 6 histone marks, DNA methylation and the transcriptome, in subjects with a history of ELA and controls. We first uncovered, in the healthy brain, previously unknown interactions among epigenetic layers, in particular related to non-CG methylation in the CAC context. We then showed that ELA associates with methylomic changes that are as frequent in the CAC as in the canonical CG context, while these two forms of plasticity occur in sharply distinct genomic regions, features, and chromatin states. Combining these multiple data indicated that immune-related and small GTPase signaling pathways are most consistently impaired in the amygdala of ELA individuals. Overall, this work provides new insights into epigenetic brain regulation as a function of early-life experience.

## Introduction

Early-life adversity (ELA), including sexual and physical abuse, as well as other forms of child maltreatment, is a major public-health problem that affects children of all socio-economic backgrounds^1^. ELA is a strong predictor of increased lifetime risk of negative mental health outcomes, including depressive disorders^2^. Among other findings, a large number of studies suggest an association between ELA and morphological and functional changes in the amygdala^3^, a brain structure critically involved in emotional regulation^4^. It is possible, thus, that amygdala changes observed in individuals who experienced ELA may contribute to increase risk of psychopathology.

The amygdala is composed of inter-connected nuclei, among which the basal and lateral sub-divisions are responsible for receiving and integrating external information. In turn, these nuclei innervate the central amygdala, the primary nucleus projecting outside the amygdalar complex to mediate behavioural outputs^4^. While specific properties of these nuclei remain difficult to assess in humans, animal studies indicate that the basal and lateral sub-divisions exhibit differential responsivity to stress, in particular as a function of the developmental timing of exposure (adolescence versus adulthood)^5,6^. Here, we focused on homogeneous, carefully dissected tissue from the human lateral amygdala.

Childhood is a sensitive period during which the brain is more responsive to the effect of life experiences^7^. Proper emotional development is contingent on the availability of a supportive caregiver, with whom children develop secure attachments^8^. On the other hand, ELA signals an unreliable environment that triggers adaptive responses, and deprives the organism from essential experience. A growing body of evidence now supports the hypothesis that epigenetic mechanisms play a major role in the persistent impact of ELA on gene expression and behaviour^9^. While DNA methylation has received considerable attention, available data also points towards histone modifications as another critical and possibly interacting factor^9^. Therefore, in this study we conducted a comprehensive characterization of epigenetic changes occurring in individuals with a history of severe ELA, and carried out genome-wide investigations of multiple epigenetic layers, and their cross-talk. Using post-mortem brain tissue from a well-defined cohort of depressed individuals with histories of ELA, and controls with no such history, we characterized 6 histone marks, DNA methylation, as well as their final endpoint at gene expression level.

We first generated data for six histone modifications: H3K4me1, H3K4me3, H3K27ac, H3K36me3, H3K9me3, and H3K27me3^10^, using chromatin-immunoprecipitation sequencing (ChIP-Seq). This allowed us to create high-resolution maps for each mark, and to define chromatin states throughout the epigenome. In parallel, we characterized DNA methylation using Whole-Genome Bisulfite Sequencing (WGBS). While previous studies in psychiatry focused on the canonical form of DNA methylation that occurs at CG dinucleotides (mCG), here we investigated both CG and non-CG contexts. Indeed, recent data has shown that non-CG methylation is not restricted to stem cells, and can be detected in brain tissue at even higher levels^11^. Available evidence also indicates that it progressively accumulates, preferentially in neurons, during the first decade of life^12,13^, a period when ELA typically occurs. Thus, we postulated that changes in non-CG methylation might contribute to life-long consequences of ELA, and focused in particular on the CAC context, where non-CG methylation is most abundant. Our results indicated that ELA leaves distinct, albeit equally frequent, traces at CG and CAC sites. Further, analyses of all epigenetic layers and the transcriptome converged to identify immune system processes and small GTPases as critical pathways associated with ELA. Altogether, these data uncovered previously unforeseen sources of epigenetic and transcriptomic plasticity, which may likely contribute to the severe and lifelong impact of ELA on behavioural regulation, and the risk of depression.

## Results

### Histone landscapes

Six histone modifications were assessed in depressed subjects with histories of ELA (n=21) and healthy controls (C) with no such history (n=17; Supplementary Tables1,2). Following the International Human Epigenome Consortium (IHEC) procedures, we achieved >60 and >30 million reads for broad (H3K4me1, H3K36me3, H3K27me3 and H3K9me3) and narrow (H3K27ac and H3K4me3) marks, respectively (4.0 billion reads total; Fig.S1a, Supplementary Table3). Quality controls confirmed that all samples for the 2 narrow marks showed relative and normalized strand cross correlations greater than 0.8 and 1.05 (Fig.S1b), respectively, according to expectations^14^. Relative to genes (Fig.1a,c), reads obtained for H3K27ac, H3K4me3 and H3K4me1 were strongly enriched around Transcription Start Sites (TSS), while H3K27me3 and H3K36me3 showed antagonistic distributions, consistent with patterns seen in other tissues^10^. Samples clustered by histone mark, with a strong distinction between activating and repressive marks (Fig.1b). To investigate tissue specificity of our dataset, we compared it with data from other brain regions and blood tissue (Fig.S2). For each modification, we observed higher correlations among amygdalar samples (r=0.75-0.92 across the 6 marks) than when compared with samples from other brain regions (r=0.51-0.81), and even lower correlations with blood mononuclear cells (r=0.35-0.64), consistent with the role of histones in tissue identity.

**Figure 1.**
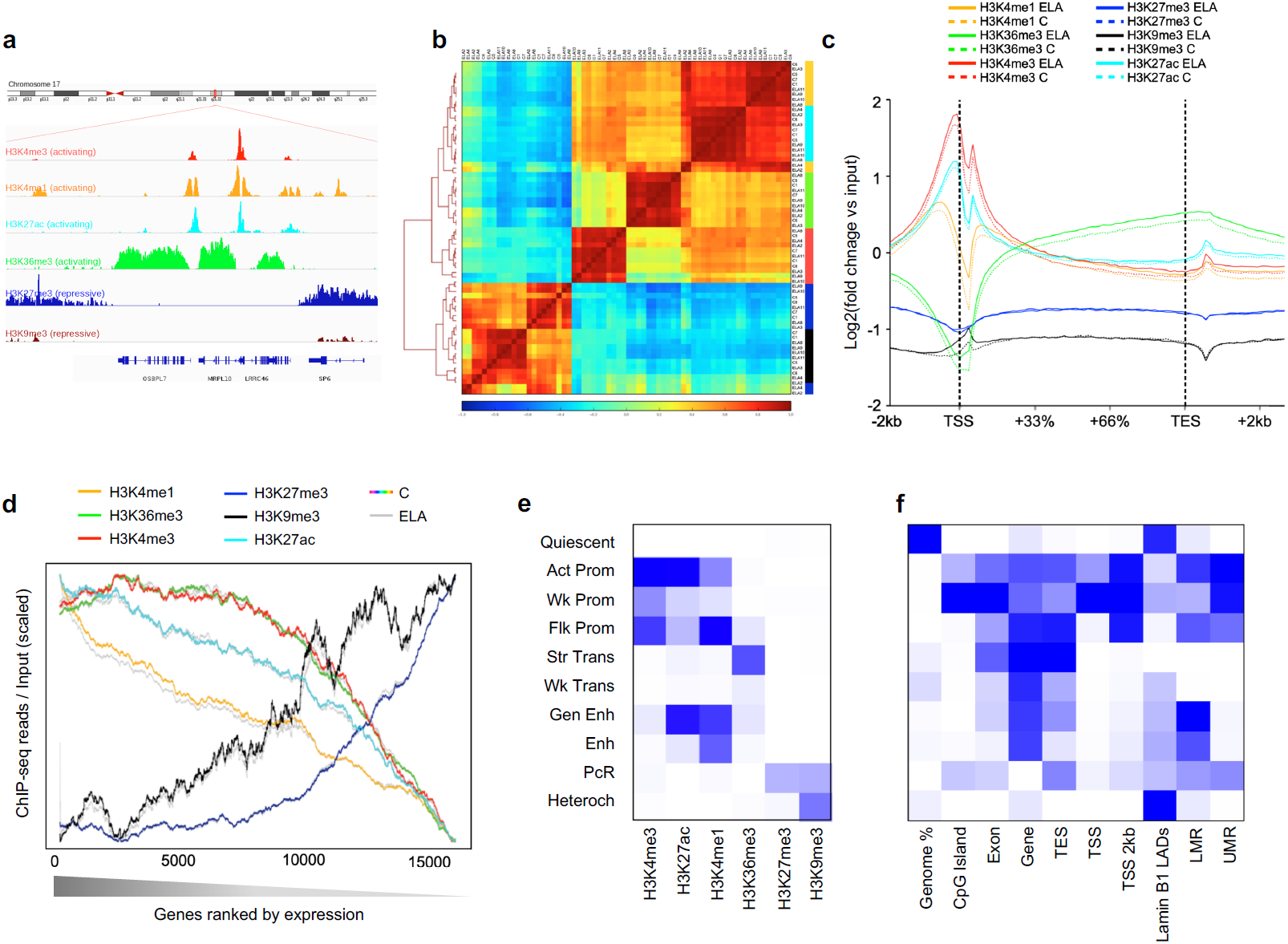
Characterization of 6 histone post-translational modifications in the human brain lateral amygdala. **(a)** Snapshots of typical ChIP-seq read distribution for the six histone marks. **(b)** Unsupervised hierarchical clustering using Pearson correlations for all marks. Correlations were computed using read number per 10 kb-bins across the whole genome, and normalized to input and library size. Note the expected separation between activating (H3K27ac, H3K36me3, H3K4me1, H3K4me3) and repressive (H3K27me3, H3K9me3) marks. **(c)** Average enrichment over input of ChIP-seq reads across all gene bodies and their flanking regions (+/- 2 kilobases, kb) in the human genome, for each histone mark. Note the expected biphasic distribution of reads around the Transcription Start Site (TSS) for H3K27ac, H3K4me3 and H3K4me1. No significant differences were observed for any mark across C and ELA groups (2-way Repeated Measures ANOVA, group effects: H3K4me1, p=0.89; H3K36me3, p=0.87; H3K4me3, p=0.64; H3K27me3, p=0.35; H3K9me3, p=0.88; H3K27ac, p=0.86). Averages for the healthy controls group (C) are shown as dashed lines, while averages for the early-life adversity group (ELA) are shown as solid lines. TES, Transcription End Site. **(d)** Average enrichment of reads over gene bodies (for H3K27me3, H3K36me3, H3K4me1 and H3K9me3) or TSS +/- 1kb (for H3K27ac and H3K4me3) for all genes ranked from most highly (left) to least (right) expressed. Strongly significant effects of gene ranking on ChIP-Seq reads were observed for all marks (p<0.0001). Again, no difference was observed as a function of ELA for any group (2-way Repeated Measures ANOVA, group effects: H3K4me1, p=0.66; H3K36me3, p=0.67; H3K4me3, p=0.98; H3K27me3, p=0.31; H3K9me3, p=0.74; H3K27ac, p=0.48). **(e)** ChromHMM emission parameters (see main text and *Methods*) for the 10-state model of chromatin generated using data from the 6 histone marks, at a resolution of 200bp, as described previously^16^. Maps of chromatin states have already been characterized in other brain regions (e.g. cingulate cortex, caudate nucleus, substantia nigra^63^) but, to our knowledge, not in the amygdala. **(f)** Intersections of chromatin states with gene features (from RefSeq) and methylomic features (lowly methylated and unmethylated regions, LMR and UMR, defined using methylseekR; see *Methods*) were computed using chromHMM’s OverlapEnrichment function. As expected, CpG-dense UMRs mostly overlapped with Promoter chromatin states (Fig.1f and Fig.S10a), while LMRs associated with more diverse chromatin states, including Enhancers (Fig.1f and Fig.S10a), consistent with their role as distant regulatory sites^31^. Chromatin states: Act-Prom, active promoter; Wk-Prom, weak promoter; Flk-Prom, flanking promoter; Str-Trans, strong transcription; Wk-Trans, weak transcription; Str-Enh, strong enhancer; Enh, enhancer; PcR, polycomb repressed; Heterochr, heterochromatin.

We next investigated relationships between histones and gene expression (Fig.1d). As expected, we observed activating functions for H3K27ac, H3K4me1, H3K36me3 and H3K4me3, and repressive functions for H3K27me3 and H3K9me3. Distinct correlative profiles were found between marks along the spectrum of gene expression, indicating that multiple marks likely better predict gene expression than individual ones. Comparisons between ELA and C groups found no significant overall differences in terms of read distribution (Fig.1c) or relationship to gene expression (Fig.1d), indicating that ELA does not globally reconfigure amygdalar histone landscapes.

Considering that different combinations of histone modifications define so-called ‘chromatin states’, we then conducted an integrative analysis of all marks using ChromHMM^15^. Maps of chromatin states were generated as described previously^16^, with each state corresponding to a distinct combination of individual marks. This unbiased approach defined a consensus map (corresponding to regions showing ≥50% agreement across all samples; Fig.1e, Fig.S3a-d and *Methods*) consistent with studies in the brain and other tissues: for example^16–18^, regions defined by H3K27ac and H3K4me1, or by H3K36me3, corresponded to known enhancers (Gen Enh and Enh) and transcribed regions (Str Trans and Wk Trans), respectively (Fig.S3e-f)^19^. Compared with known genomic features (Fig.1f), this map showed expected enrichments of promoter chromatin states (Act, Wk, or Flk Prom) at transcription start sites and CpG islands, and of transcription states (Str Trans and Wk Trans) within genes. Finally, the chromatin states exhibit expected correlations with gene expression (Fig.S4). As detailed below, these maps allowed us to characterize cross-talks between chromatin and DNA methylation, and differences between groups.

### CG and non-CG methylation patterns

We used WGBS to characterize the amygdala methylome. Rates of bisulfite conversion, sequencing depth and library diversity met IHEC standards and were similar across groups (Fig.S5a-d). In this large dataset, >13 million CGs showed an average coverage ≥5 in the cohort (Fig.S5e), which favourably compares with recent human brain studies in terms of sample size^20^ or CGs covered^21,22^.

Because non-CG methylation is enriched in mammalian brains^11,23^, we first computed average genome-wide levels of methylation in multiple cytosine contexts. Focusing on 3-letter contexts (Fig.2a), we observed that, as expected, methylation levels were highly variable among the 16 possibilities (2-way ANOVA; context effect: [F(15,540)=196283; p<0.0001]), with much higher methylation levels in the CGN contexts than in the 12 non-CG contexts. Of note, no difference was found in overall methylation between groups ([F(1,36)=0.12; p=0.73]), indicating that ELA does not associate with a global dysregulation of the methylome. Among non-CG contexts, as previously described by others in mice^24^ or humans^25,26^ (Fig.S6a-b), methylation levels were highest at CACs (4.1±0.1%), followed by a group of contexts between 1.8 to 1.1% (CTC, CAG, CAT, and CAA), and remaining ones below 0.4%. Considering that methylation at CA^27^ or CAC^28^ sites may have specific functions in the brain, and because CAC methylation (hereafter mCAC) was most abundant, we focused on this context.

**Figure 2.**
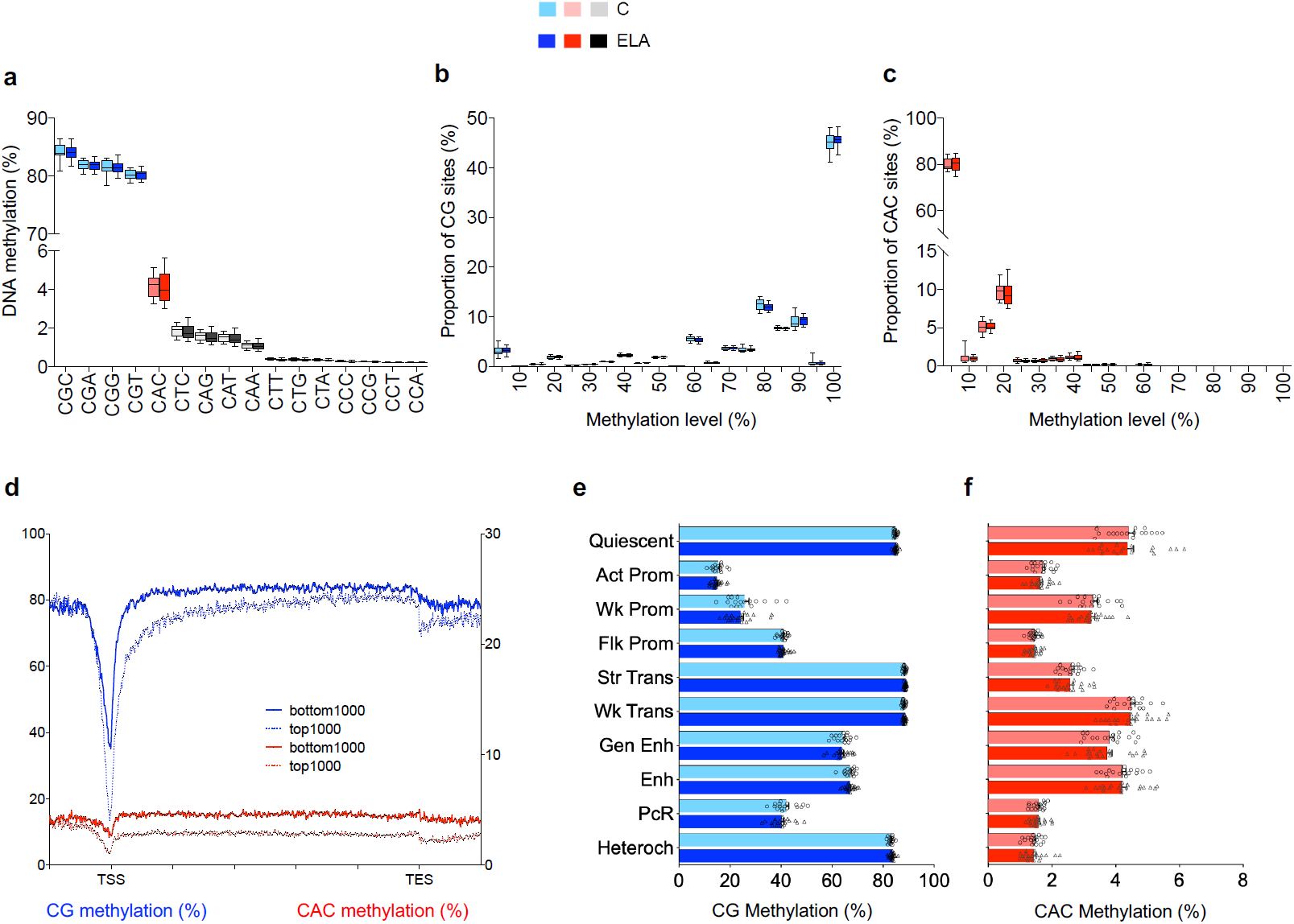
Characterization of non-CG methylation in the human brain lateral amygdala. **(a)** Average genome-wide levels of DNA methylation were measured among the sixteen 3-letter cytosine contexts (CHH, where H stands for A, C, and T) in the human brain lateral amygdala, using whole-genome bisulfite sequencing. While highest DNA methylation levels were observed in the 4 CGN contexts (where N stands for any base; CGC: 84.1±0.2%; CGA: 81.9±0.1%; CGC: 81.4±0.2%, CGT: 80.2±0.1%; mean±sem in the whole cohort), detectable non-CG methylation was also observed in CHN context, most notably at CAC sites (4.1±0.1% in combined control, C, and early-life adversity, ELA, groups), with no detectable differences between groups for any context (2-way Repeated Measures ANOVA; group effect: [F(1,36)=0.12; p=0.73]). **(b)** DNA methylation in the CG context mostly corresponded to sites highly methylated. In contrast, as previously described in the mouse hippocampus^29^, most CAC sites were unmethylated **(c)**, with only a minority of sites showing low methylation levels, between 10 and 20%. This likely reflects the fact that non-CG methylation does not occur in all cell types, and is notably enriched in neuronal cells and, to a lesser extent, in glial cells^11^. In the CG or CAC contexts (2-way ANOVA; group effect: CG, [F(1,720)=5.0E-11; p>0.99]; CAC, [F(1,36)=0; p>0.99]), ELA did not associate with any significant change in these global distributions. Box plots show median and interquartile range, with whiskers representing minimum and maximum values. **(d)** In both contexts, patterns of DNA methylation along gene bodies showed the expected anti-correlation with gene expression, as shown here comparing the 1000 most highly (top1000) or lowly (bottom1000) expressed genes, consistent with previous rodent data. TSS, Transcription Start Site; TES, Transcription End Site. In the CG **(e)** or CAC **(f)** contexts, no difference in DNA methylation levels was observed between C and ELA groups for any chromatin state. We observed, however, dissociations in the relationship of DNA methylation and histone marks across the CG and CAC contexts (see main text). Values are mean±sem. Chromatin states: Act-Prom, active promoter; Wk-Prom, weak promoter; Flk-Prom, flanking promoter; Str-Trans, strong transcription; Wk-Trans, weak transcription; Str-Enh, strong enhancer; Enh, enhancer; PcR, polycomb repressed; Heterochr, heterochromatin.

We first compared mCG and mCAC. While CG sites were highly methylated, CAC (Fig.2b-c) or other non-CG (Fig.S6c) sites were mostly unmethylated, with a minority of them showing methylation levels between 10 to 20%, consistent with mouse data^29^. Regarding distinct genomic features and chromosomal location, we confirmed that: (i) while mCG is lower within promoters, this effect is much less pronounced for mCAC (Fig.S7a)^11^; (ii) compared with CGs^30^, depletion of methylation from pericentromeric regions is even stronger at CACs, and (iii) as expected, methylation levels were extremely low in both contexts in the mitochondrial genome (Fig.S7b). We also confronted methylation data with gene expression, regardless of group status, and found the expected anti-correlation in both contexts (Fig.2d; CG: [F(1,37)=557; p=6.7E-24]; CAC: [F(1,37)=3283; p=9.7E-38]). Because CAC sites, in contrast with CGs, are asymmetric on the two DNA strands, we wondered whether this anti-correlation would be different when contrasting gene expression with mCAC levels on its sense or antisense strand (Fig.S8). No difference was found, indicating that gene expression is predicted to the same extent by mCAC on either strand, at least for the coverage achieved here. Finally, we used methylseekR to characterize, to our knowledge for the first time in the human brain, active regulatory sites defined as unmethylated (UMR) and lowly methylated (LMR) regions (Fig.S9a-e). As observed in other tissues, CG-dense UMRs mostly overlapped with CpG islands and promoter chromatin states (Fig.1f, Fig.S9d, Fig.S10a), while LMRs associated with more diverse states (including Enhancers; Fig.S1f, Fig.S10a), consistent with their role as distant regulatory sites^31^. Among each LMR and UMR category, significant variations in levels of mCG or mCAC were observed across various chromatin states (Fig.S9f). Regarding individual histone marks at LMR and UMR, we further documented specific associations, including patterns of depletion and enrichment specific to UMR shores not characterized previously (see Fig.S10b-g for details). Overall, these differences and similarities between mCG and mCAC extends on results obtained previously in smaller cohorts of mouse or human samples^29,32^.

Regarding histone modifications, while mechanisms mediating their interactions with mCG have been documented, no data is available to describe such relationship for non-CG contexts. To address this gap, we confronted our consensus model of chromatin states with DNA methylation (Fig.2e-f). Levels of mCG ([F(9,324)=5127; p<0.0001]) and mCAC ([F(9,324)=910.7; p<0.0001]) strongly differed between states, unravelling previously uncharacterized patterns. First, lowest levels of mCG were found in the 3 promoter states (Fig.2e), corresponding to a strong anti-correlation between DNA methylation and both forms of H3K4me1,3 methylation, consistent with previous findings in other cell types^33^. Accordingly, these 3 promoter states were defined (Fig.1e) by high levels of H3K4me3 in combination with either: (i) high H3K4me1 (Flanking Promoter, Flk Prom; p<0.0001 for every post-hoc comparison, except against the Polycomb repressed state, PcR); (ii) high H3K27ac (Active Promoter, Act-Prom; p<0.0001 for every comparison against other states), or (iii) intermediate levels of both H3K27ac and H3K4me1 (Weak Promoter, Wk-Prom; p<0.0001 against other states). In contrast, among these 3 promoter states mCAC was particularly enriched in Wk-Prom regions (p<0.0001 against Act-Prom and Flk-Prom; Fig.2f, Fig.S3d). Second, mCG was abundant in transcribed regions defined by either intermediate (Weak Transcription, Wk-Trans) or high (Strong Transcription, Str-Trans) H3K36me3. By contrast, mCAC was selectively decreased in the Str-Trans state (p<0.0001 against Wk-Trans). Third, while mCG levels were high in heterechromatin (Heteroch, defined by high H3K9me3), consistent with its role in chromatin condensation, mCAC appeared depleted from these regions (p<0.0001 for every comparison against other states, except PcR and Flk-Prom). These results indicate that interactions between DNA methylation, histones, and chromatin strikingly differ across mCG and mCAC, possibly as a result of brain-specific epigenetic processes in the latter 3-letter context^32^. Finally, as expected, ELA did not associate with a global disruption of this cross-talk, as no changes in genome-wide levels of mCG ([F(1,36)=0.36; p=0.55]) or mCAC ([F(1,36)=0.07; p=0.80]) were observed across C and ELA groups for any state.

### Changes in histone marks and chromatin states as a function of ELA

We investigated local histone adaptations in ELA subjects using diffReps^34^. A total of 5126 differential sites (DS) were identified across the 6 marks (Fig.3a-b, Fig.S11, Supplementary Table4) using consensus significance thresholds^35^ (p<10^−4^, FDR-q<0.1). Interestingly, H3K27ac contributed to 30% of all DS, suggesting a prominent role of this mark. Annotation to genomic features revealed distinct distributions of DS across marks (df=25, *χ*^*2*^ =1244, p<0.001; Fig.S12a): H3K4me1- and H3K4me3-DS were equally found in promoter regions and gene bodies, while H3K36me3- and H3K27ac-DS were highly gene-body enriched, and H3K27me3- and H3K9me3-DS found in intergenic/gene desert regions. Sites showing enrichment (up-DS) or depletion (down-DS) of reads in ELA subjects were found for each mark, with an increased proportion of down-DS associated within H3K4me1, H3K4me3, H3K36me3 and H3K27me3 changes (Fig.S12b).

**Figure 3.**
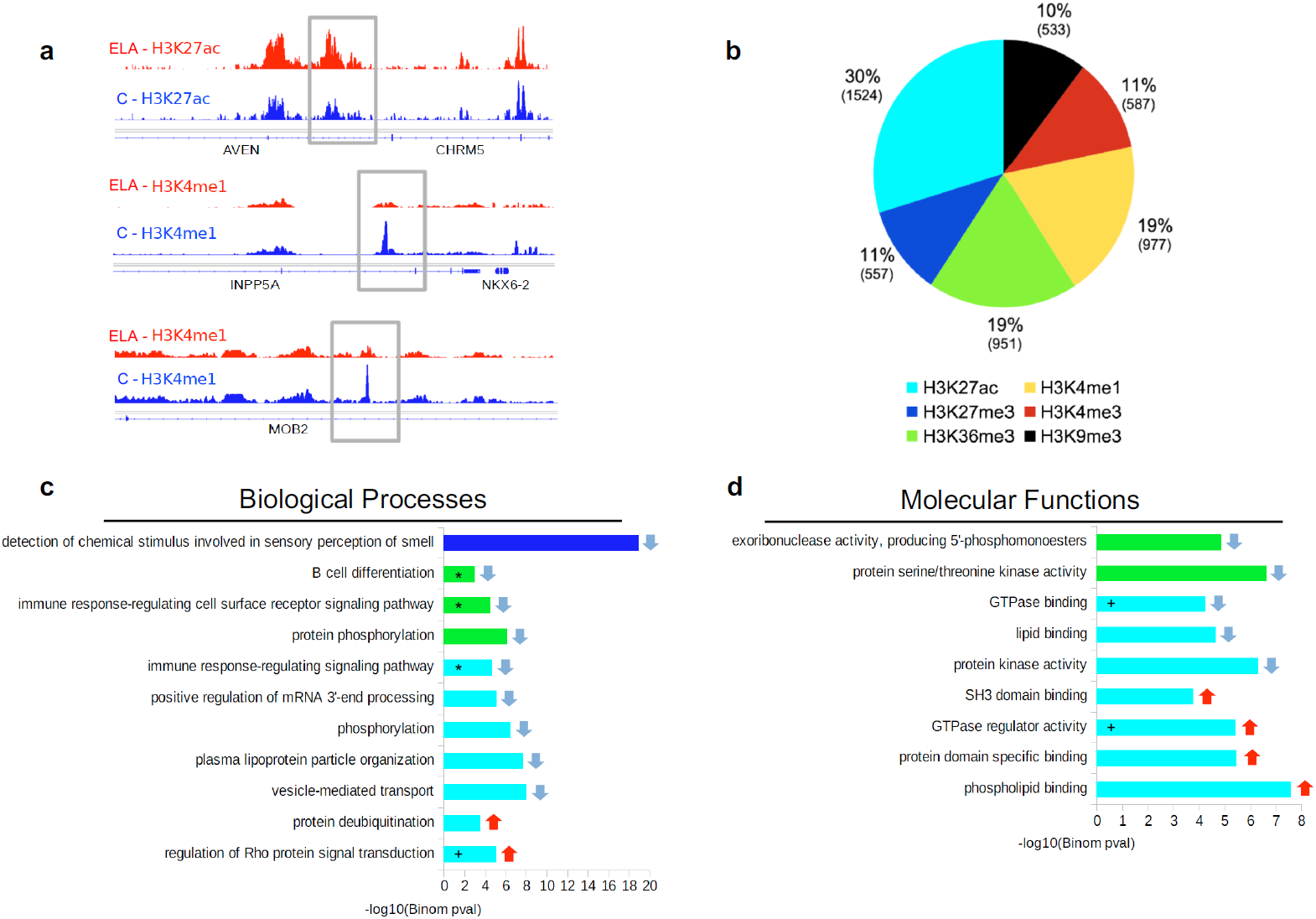
Analysis of genomic sites showing differential enrichment for individual histone marks in subjects with a history of early-life adversity (ELA). **(a)** Representation of three top Differential Sites (DS), identified using diffReps^34^. ELA are shown in red, healthy controls (C) are shown in blue. Grey rectangles delineate the coordinates of each DS. **(b)** Relative proportion of DS contributed by each histone mark. Percentages of total number of DS, and absolute number of DS (in brackets) are shown for each mark. Both depletion- and enrichment-DS were observed for each of the 6 marks (Fig.S12b). Among genes most strongly affected (Supplementary Table4), several have been previously associated with psychopathology, such as QKI (H3K27ac top hit)^40,68^ or HTR1A (H3K4me3 top hit)^69^. **(c-d)** Top five most significant non-redundant gene ontology “Biological Processes” **(c)** or “Molecular Functions” **(d)** terms enriched for each histone mark DS, as identified by GREAT^36^ using hypergeometric and binomial testing (fold change≥1.5 and FDR-q≤0.1 for both tests). Surprisingly, the single most significant result implicated epigenetic dysregulation of odor perception in ELA subjects (consistent with recent clinical studies^70^), while immune processes (indicated by *), and small GTPases (+) were consistently found affected across different marks. Negative logarithmic p-values are shown for binomial testing. Color indicates histone mark concerned, arrows indicate direction of event: terms associated with depletion- (down arrow) or enrichment-DS (up arrow).

We then used GREAT (Supplementary Table5), a tool that maps regulatory elements to genes based on proximity, to test whether ELA subjects had histone modifications affecting genes in specific pathways^36^. We performed this GO analysis on each mark and found significant enrichments for 3 of them (Fig.3c-d). Importantly, overlaps between enriched GO terms were observed across these 3 marks: notably, terms related to immune processes, as well as small GTPases and Integrin signaling (Fig.S12c) were enriched for H3K36me3- and H3K27ac-DS, suggesting these pathways may play a significant role in ELA.

To strengthen these findings, a complementary analysis was conducted using chromatin state maps^15^. First, we identified genomic regions where a state transition (ST; n=61,922) occurred between groups (Supplementary Table6). Across the 90 possible ST in our 10-state model, only 56 were observed, with a high proportion (50.2%; * in Fig.4a) involving regions in quiescent (Quies), Wk-Trans or Enh states in the C group that mostly turned into Quies, Str-Trans, Wk-Trans, and Heteroch states in the ELA group. Furthermore, 17% and 59% of ST occurred in regions within 3kb of a promoter or in gene bodies (Fig.4b), respectively, suggesting that ELA-associated changes affected selected chromatin states, and mostly occurred within genes.

**Figure 4.**
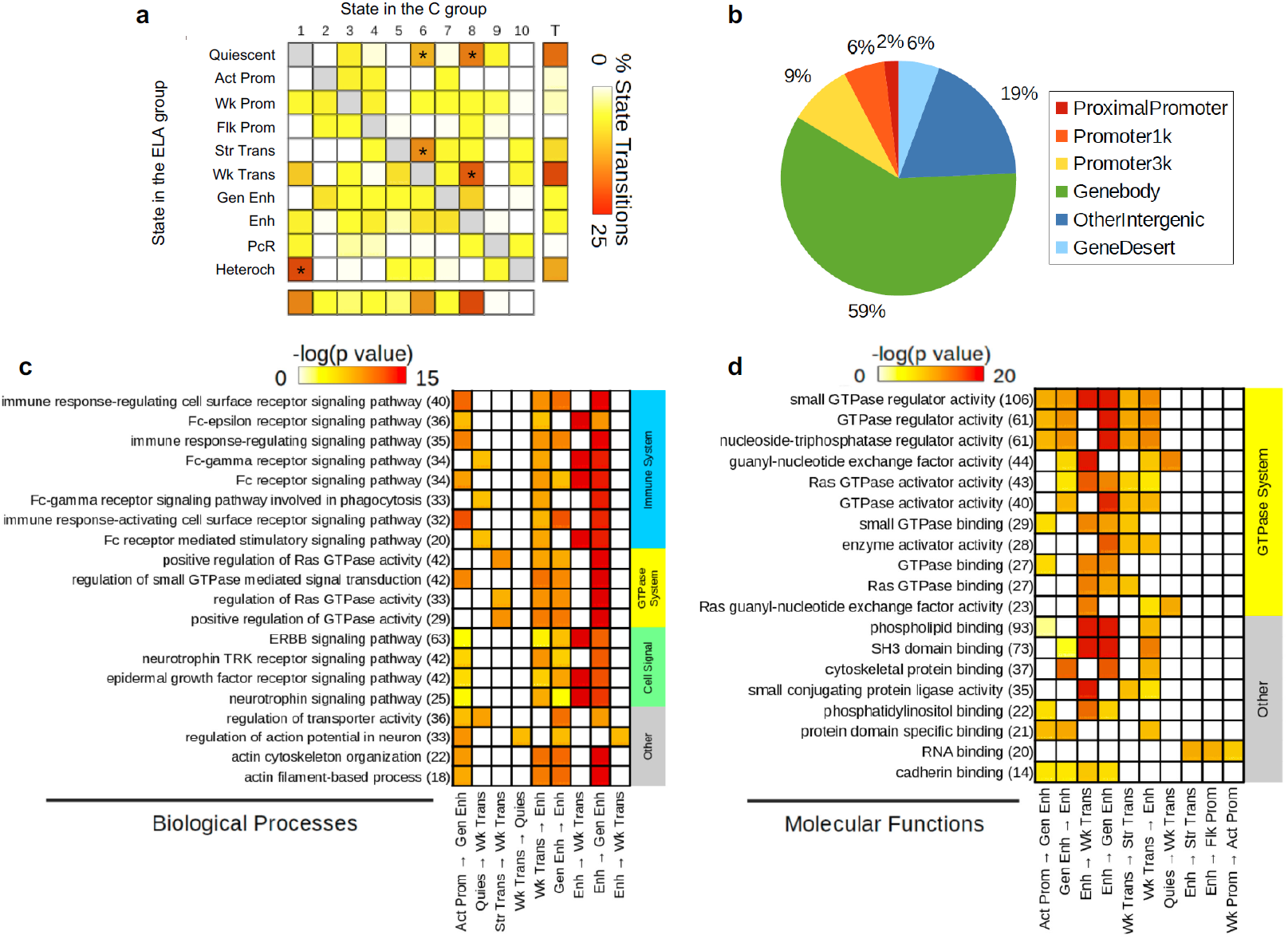
Analysis of genomic sites showing a switch between chromatin states as a function of early-life adversity (ELA). **(a)** Percentage of each State Transition (ST) type relative to the total number of transitions. For the healthy control (C) versus ELA group comparison, the cumulative percentages of ST from a specific state to any other state are shown in the “Total” and “T” rows/columns. * indicates most frequent STs (see main text). **(b)** Distribution of ST localizations relative to genomic features, assessed using region_analysis^34^ (see *Methods*; pericentromeric and subtelomeric categories not shown). **(c-d)** Gene ontology “Biological Processes” **(c)** or “Molecular Functions” **(d)** terms significantly associated with at least three types of ST. Terms are grouped based on overall system involved, and ranked by co-occurrence score (in parentheses after each term), which reflects both the significance of GO terms and their recurrence across multiple ST (see main text and^35^). Individual binomial p-values for each type of ST and each term are shown by color gradient. Each term also passed hypergeometric testing. Immune-related and small GTPase terms were most strongly affected, across multiple ST. Of note, a complementary GREAT pathway analysis using MSigDB further strengthened these findings by revealing recurrent enrichment of the integrin signalling pathway (across six types of ST, as well as for H3K27ac down-DS; see Fig.S12c), which is known to interact extensively with small GTPases^71^.

We next investigated GO enrichment of ST using GREAT (Fig.4c-d, Supplementary Table7) and a co-occurrence score reflecting both the significance of GO terms and their recurrence across multiple ST^35^. Importantly, biological processes (Fig.4c) with highest co-occurrence scores were similar to those found from the GO analysis of individual histone marks, and clustered in two main categories: immune system, and small GTPases. These terms were significant for ST involving transcription, quiescent and enhancer states. Regarding molecular functions (Fig.4d), most enriched categories were related to GTPases, and involved the same types of ST. Therefore, analyses of individual histone marks and chromatin state converged to suggest impairments in similar GO pathways.

### Differential DNA methylation in ELA

We next sought to identify changes in DNA methylation. As mCG and mCAC were very different, and considering data suggesting possible mCAC-specific processes^28^, we used BSmooth^37^ to identify DMRs separately in each context, with strictly similar parameters (see *Methods*). DMRs were defined as regions of ≥5 clustered cytosines that each exhibited a significant group difference in methylation (p<0.001). Also, because age and sex are known to affect DNA methylation^38,39^, generalized linear models were computed for each DMR, and only those that remained significant when taking these 2 covariates into account were kept for downstream analyses. Surprisingly, we found that as many DMRs could be identified in the CAC (n=840) as in the canonical CG (n=795) context, suggesting that cytosines in the CAC context may represent a significant form of plasticity.

While both types of DMRs were similarly abundant and distributed throughout the genome (Fig.5a-b), they nevertheless showed striking differences. Compared with CG-DMRs, CAC-DMRs were composed of slightly fewer cytosines (Fig.S13a, p=2.9E-04) and smaller (Fig.S13b, p<2.2E-16). CG-DMRs also affected sites showing a wide range of methylation levels, while CAC-DMRs were located in lowly methylated regions (Fig.5c-d), consistent with genome-wide lower mCAC levels. In addition, the magnitude of methylation changes detected in the ELA group were less pronounced in the CAC context, with smaller % changes (p<2.2E-16; Fig.5e-f, Fig.S13c) and areaStat values (the statistical strength of DMRs^37^; p=5.8E-08, Fig.S13d). Further strengthening differences between the 2 contexts, CG- and CAC-DMRs showed no genomic overlap (Fig.S13e) and very distinct distributions among UMR and LMR features (Fig.5g). Finally, when considered collectively, genomic regions were CG-DMRs were identified as a function of ELA showed no group difference in the CAC context (and vice versa, see Fig.5h), indicating that ELA-related processes do not simultaneously affect both cytosine contexts.

**Figure 5.**
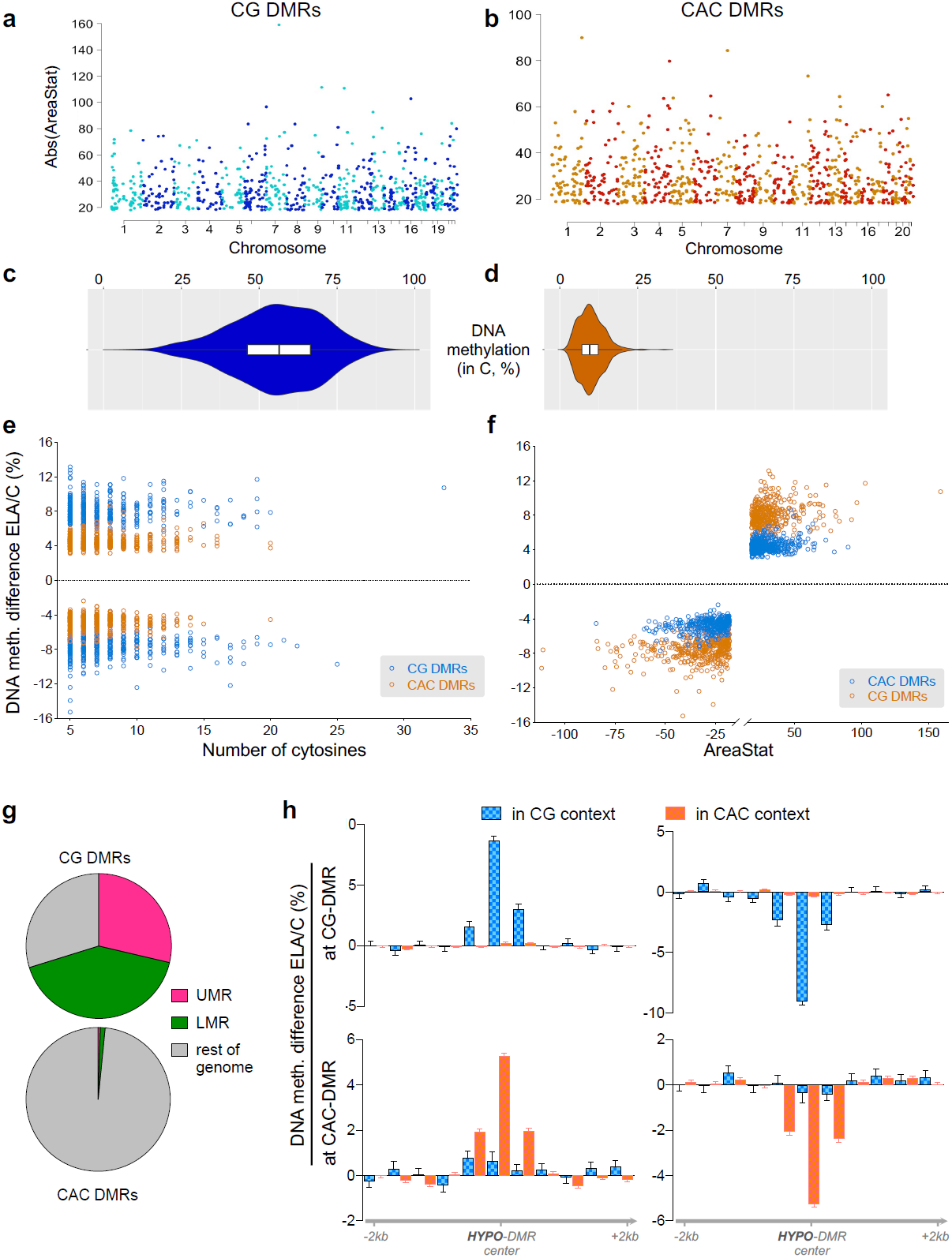
Differential DNA methylation in the CG and CAC contexts in subjects with a history of early-life adversity (ELA). **(a-b)** Manhattan plots of differentially methylated regions (DMR) identified using the BSmooth algorithm in the CG and CAC contexts, comparing control (C) and ELA groups. DMRs were identified separately in each context using the BSmooth algorithm^37^, with strictly similar parameters (see *Methods*). They were defined as regions of ≥5 clustered cytosines that each exhibited a significant difference in methylation (p<0.001) and an absolute methylation difference ≥1% between groups. Surprisingly, as many DMRs were identified in the CAC context (n=866) as in the canonical CG context (n=878). **(c-d)** Methylation abundance in the C group in regions where DMRs were identified in the CG and CAC contexts. CG-DMRs affected genomic sites showing a wide range of methylation levels (mean±sem=55.3±0.5%), while CAC-DMRs occurred in lowly methylated regions (mean±sem=10.0±0.1%), resulting in significantly different distributions (Mann-Whitney U=686; p<0.0001). Box plots show median and interquartile range (IQR), with whiskers representing 1.5 × IQR. **(e)** DNA methylation differences observed in ELA subjects compared to the C group in CG and CAC DMRs, as a function of the number of cytosines composing each DMR. **(f)** DNA methylation differences observed in ELA subjects compared to the C group in CG- and CAC-DMRs, as a function of areaStat values, a measure of the statistical significance of each DMR implemented by BSmooth. **(g)** Distinct distributions (χ^2^=884.3, df=2, p<1E-15) of CG- and CAC-DMRs among lowly methylated (LMR) and unmethylated (UMR) regions**. (h)** DNA methylation in CG and CAC contexts at DMRs. No average difference in mCAC levels were observed among C and ELA groups at CG-DMRs (upper panels) that showed either increased (HYPER-DMR) or decreased (HYPO-DMR) levels of methylation in ELA subjects (and vice versa for CAC-DMRs, lower panels).

We next characterized genomic features where DMRs occurred, and observed that their distribution again strikingly differed (p<2.2E-16; Fig.6a-b, Supplementary Table8): CG-DMRs were located in promoters (38.5% in proximal promoter, promoter1k and promoter3k) and gene bodies (35.4%), while CAC-DMRs were mostly in gene bodies (53%) and intergenic regions (28.1%). Second, we characterized histone modifications around DMRs (Fig.6c-d, Fig.S15a-f): CG-DMRs were enriched with H3K4me1, H3K4me3 and H3K27ac (Fig.6c), coherent with our observations that these histone marks (Fig.1e) and DMRs (Fig.6a) preferentially located at promoters. In sharp contrast, the 2 main features characterizing CAC-DMRs were an enrichment in H3K36me3 and a depletion in H3K9me3 (Fig.6d, Fig.S15e-f). These differences were further supported by the analysis of chromatin states (p<2.2E-16; Fig.6e, Fig.S15g, Supplementary Table9). CAC-DMRs were largely absent from promoter (Act-Prom, Flk-Prom, and Wk-Prom) and enhancer (Str-Enh and Enh) states that were all defined, to varying degrees, by the 3 marks that primarily characterize CG-DMRs: H3K4me1, H3K4me3 and H3K27ac (Fig.1e). In addition, CAC-DMRs were (i) enriched in the Wk-Trans state, defined by the presence of H3K36me3, and (ii) depleted from the 2 states (PcR, Heteroch) characterized by H3K9me3.

**Figure 6.**
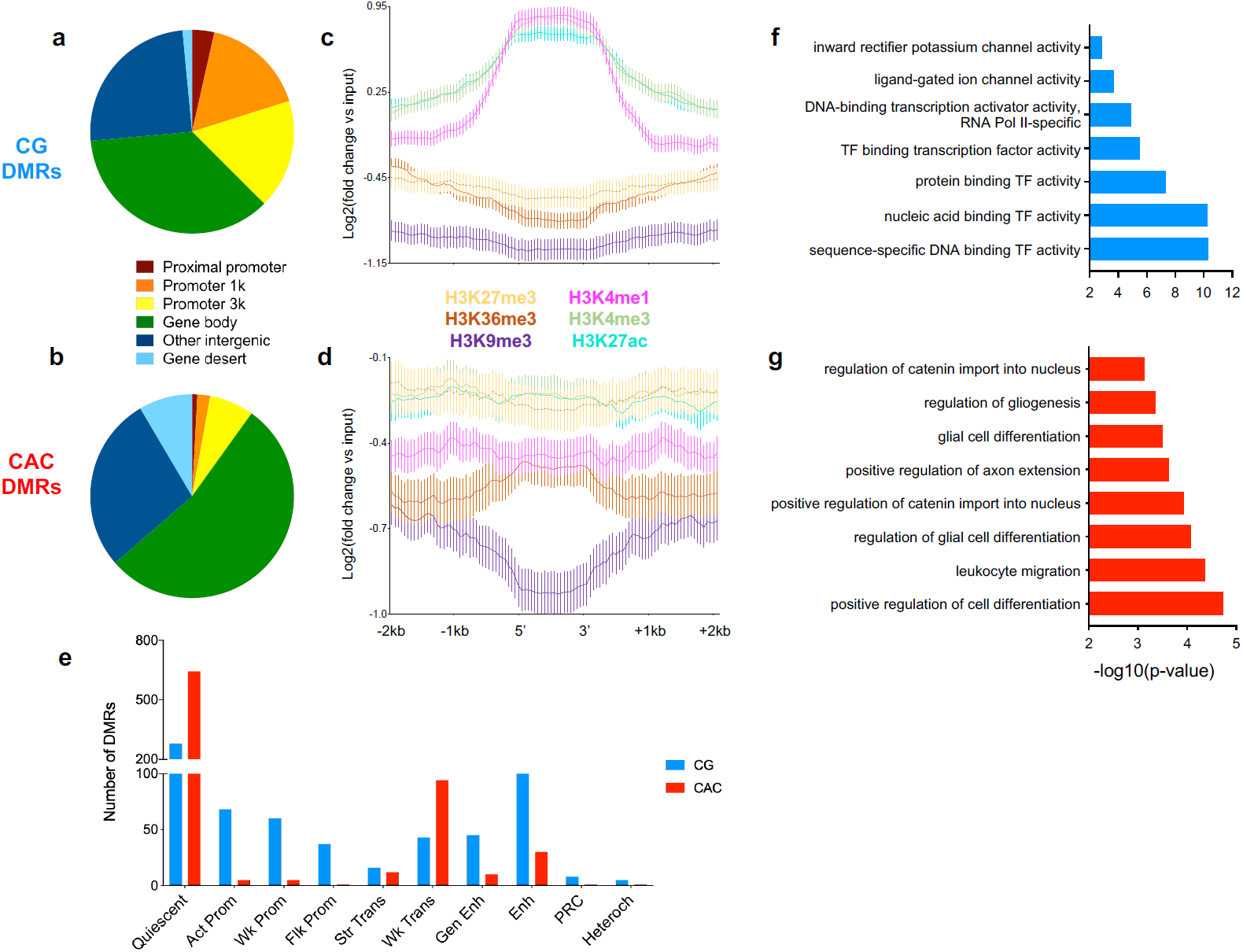
Individual histone marks and global chromatin states defining genomic regions where early-life adversity (ELA) associated with differential DNA methylation. **(a-b)** Localization of differentially methylated regions (DMR) in genomic features, identified using region_analysis^34^. Distributions were strongly different among CG and CAC contexts (χ^2^=221.2, df=6, p<2.2E-16). **(c-d)** Histone modifications measured at the level of DMRs and their flanking regions (+/- 2 kilobases, kb). Distributions were very distinct between CG- and CAC-DMRs, with significant interactions between cytosine context and cytosine position along DMRs, for each of the 6 marks (2-way Repeated Measures ANOVA interactions, p<0.0001 for all; see also Fig.S15a). Values are mean±sem. **(e)** Chromatin states found at DMRs. Similarly, CG- and CAC-DMRs occurred in very different chromatin states (χ^2^=390.4, df=9, p<2.2E-16). **(f-g)** Gene Ontology analysis of CG and CAC DMRs using GREAT^36^ (see main text).

Finally, we conducted a GREAT analysis of GO terms enriched for DMRs: CG-DMRs notably associated with terms related to the regulation of neuronal transmembrane potential (Fig.6f and Supplementary Table10), in agreement with histone results (Fig.4c), while CAC-DMRs were enriched for terms related to glial cells (Fig.6g), consistent with the immune dysregulation previously observed with histone DS and ST. Altogether, while ELA associates with similar numbers of mCG and mCAC adaptations, these 2 types of plasticity occur in genomic regions characterized by different histone marks, chromatin states, and GO categories, possibly reflecting the implication of distinct molecular mechanisms.

### Differential gene expression in ELA and combined GO analyses

Analyses of histones and DNA methylation identified GO terms consistently affected in ELA individuals. To determine how these epigenetic adaptations may ultimately modulate amygdalar function, we characterized gene expression in C and ELA groups using RNA-Sequencing. Samples with similar RNA integrity across groups were sequenced at >50 million reads/sample (Fig.S16). Quantification of gene expression was conducted using HTSeq-count^40^ and validated by an alternative pseudo-alignment approach, Kallisto^41^, generating very similar results (r=0.82, p<2.2E-16; Fig.S17a). A differential expression analysis between groups was then performed using DESeq2 (Supplementary Table11). Similar to our epigenetic analyses, we searched for patterns of global functional enrichment, using GO and Gene Set Enrichment Analysis (GSEA)^42^. Enrichment of GO categories using genes that showed nominal differential expression in the ELA group (p<0.05, n=735, Fig.7a, Supplementary Table12) identified numerous terms consistent with previous analyses at epigenetic level, including immune and small GTPase functions (Fig.7b). We also used GSEA^42^, which does not rely on an arbitrary threshold for significance, and takes directionality of gene expression changes into account. GSEA identified 163 genome-wide significant sets, among which 109 were related to immune processes and negatively correlated with ELA (Supplementary Table13, Fig.7c-d, Fig.S17b). Therefore, transcriptomic data revealed gene pathways that in part overlap with those identified using histone marks and DNA methylation.

**Figure 7.**
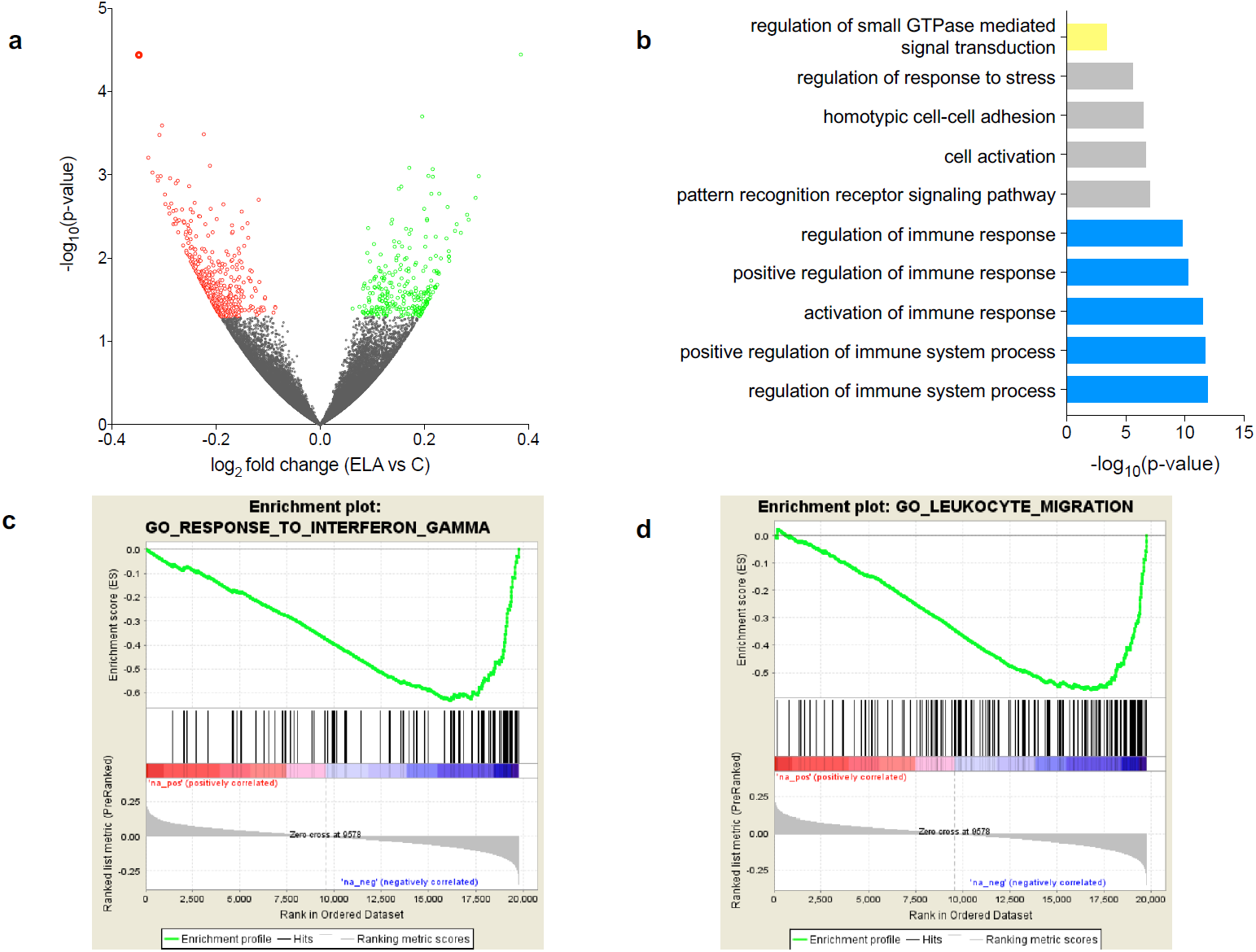
Differential gene expression in subjects with a history of early-life adversity (ELA). **(a)** Volcano Plot of RNA-Seq data showing the 261 and 474 genes that were up- (green circles) or down- (red circles) regulated in the ELA group compared with the control (C) group (nominal p-value<0.05). **(b)** Gene Ontology analysis of the 735 differentially expressed genes in the ELA group. Terms showing evidence of enrichment for differential methylation, histone profile or chromatin state are shown in yellow (small GTPase) or blue (immune processes; see also Fig.S17c). **(c-d)** Gene Set Enrichment Analysis (GSEA) of gene expression changes in ELA subjects. Genes were ranked based on log_2_ fold changes from the C versus ELA differential expression analysis (‘Ranked list metric’, in grey in lower portion of each panel). Genes with highest positive fold changes (in red, upregulated in the ELA group) are at the extreme left of the distribution, and those with lowest negative fold changes (in blue, downregulated in the ELA group) are at the extreme right. A running enrichment score (green line, upper portion of each panel) was computed for gene sets from the MSigDB curated molecular signatures database, and used to identify enriched gene sets^42^. Among the numerous gene sets related to immune function that showed evidence of genome-wide significant negative correlation with ELA (see main text, and Supplementary Table13), 2 representative gene sets are shown (with the middle portion of each panel showing vertical black lines where members of the gene set appear in the ranked list of genes): (c) ‘Interferon gamma’, and (d) ‘Leukocyte migration’. Of note, an oligodendrocyte-specific gene collection, which we recently found downregulated in the cingulate cortex of subjects with a history of ELA^40^, positively correlated with ELA in the amygdala (see Fig.S17b), suggesting opposite adaptations in this glial population between cortical and subcortical structures.

Because epigenetic and transcriptomic patterns determine and reflect cellular identity, adaptations associating with ELA in the present work may stem from changes in cellular composition of the amygdala. To explore this possibility, we deconvoluted our bulk tissue measures of gene expression and DNA methylation using BSEQ-sc^43^ and the Cibersort^44^ algorithm. For gene expression, we used as reference single-nucleus transcriptomes recently generated by our group using cortical tissue^45^. Results showed proportions of excitatory (80%) and inhibitory (20%) neuronal subtypes consistent with expectations (Fig.S14a) while, importantly, no changes in proportions of neuronal populations, microglia, astrocytes, or oligodendrocytes could be identified in these analyses (Fig.S14b-c). For DNA methylation, we used as reference single-cell non-CG methylomes recently published^46^, and found some convergence with cellular estimates generated using RNA-Sequencing (Fig.S14d). Again, estimated abundance of different classes of excitatory and inhibitory neurons across C and ELA groups were unchanged (Fig.S14e), reinforcing the hypothesis that ELA-related adaptations reflect changes in cellular phenotypes rather than abundance.

To combine analyses conducted for histones, chromatin states, DNA methylation and gene expression, we finally grouped GO terms enriched at each level to identify biological mechanisms most consistently affected (Fig.S17c). Overall, a clear pattern emerged whereby the highest number of genome-wide significant terms (n=125 GO terms) were related to immune processes, with contributions from each of the 4 types of data. Second came terms related to small GTPases, which were documented by histone modifications, chromatin states and gene expression (n=22), followed by terms related to neuronal physiology (n=19, mostly linked with neuronal excitability and sensory processing; Supplementary Table14), cellular adhesion (n=11), and the cytoskeleton (n=6). Altogether, these combined analyses defined major epigenetic and transcriptomic pathways affected by ELA in the lateral amygdala.

## Discussion

Imaging studies^3^ have consistently demonstrated that ELA associates with impaired amygdala function. Here, going beyond previous studies^9^, we conducted a comprehensive analysis of its potential molecular consequences in this brain region across multiple transcriptional and epigenetic mechanisms.

Over the last two decades, considerable evidence has associated enhanced inflammation with stress-related phenotypes such as depression, in particular based on measures of cytokines and inflammatory factors in blood samples^47^. Limited molecular data, however, is available to understand how this pro-inflammatory state may translate in the brain. Available studies focused on cortical structures of the frontal lobe, and reported conflicting results for the expression of related genes^48–50^. At histological level, while the prevailing view holds that stress-related psychopathology associates with tissue inflammation^47^, studies conducted on the amygdala showed discordant results, with lower densities of glial cells in some^51,52^ but not all^53^ studies. In the present work, integration of genome-wide data on DNA methylation, histone and gene expression found converging evidence for significant enrichment in immune-related GO terms across all molecular layers. This included decreased expression of genes encoding the complement system, Toll-like receptors, clusters of differentiation, and the major histocompatibility complex, altogether arguing for a meaningful contribution to psychopathological risk. Of note, deconvolutions of transcriptomic or methylomic data provided no indications of changes in amygdala cellular composition as a function of ELA, suggesting that reported molecular adaptations may reflect decreased activity rather than impaired recruitment or proliferation of microglial and astrocytic cells, the main immune actors in brain tissue. Altogether, our data therefore suggest that dysregulation of immune-related processes in the amygdala may play an important role in long-term consequences of ELA and in the pathophysiology of depression.

While proteins from immune pathways have been historically identified and studied in the context of immune function and associated circulating cells (eg lymphocytes, monocytes and granulocytes), a growing and significant literature now indicates that they are also largely expressed by neuronal cells, and play important role in the regulation of synaptic plasticity^54–59^. Consistently, other pathways most significantly altered in ELA subjects was related to small GTPases, a large family of GTP hydrolases that regulate synaptic structural plasticity, notably through interactions with the cytoskeleton^60^. Interestingly, the association observed for small GTPases was also accompanied by significant changes affecting GO terms related to the cytoskeleton. Overall, our findings therefore point towards altered synaptic plasticity in the lateral amygdala in relation to ELA and depression, and reveals part of underlying epigenetic mechanisms at DNA methylation and histone levels. While very few molecular studies in humans previously documented this hypothesis^61^, it strongly resonates with the wealth of human imaging and animal data that shows structural and functional plasticity in this brain region as a function of stressful experiences^4^.

Next, we conducted a detailed analysis of non-CG methylation. Over the last few years, the significance of this type of DNA methylation, and the possibility that it may fulfill biological functions, have been supported by several lines of evidence, including: (i) distinct methylation patterns shown to preferentially affect CAG sites in embryonic stem cells as opposed to CACs in neuronal and glial cells^11^, (ii) the particular abundance of non-CG methylation in genes with a higher genomic size in the human brain^27^, potentially implicated in Rett syndrome, and (iii) the specific binding of the methyl-CpG-binding domain protein MeCP2 to both mCG and mCAC in the mouse brain^28,62^. Here, we provide additional evidence reinforcing this notion, and found that mCAC exhibits peculiar interactions with the histone code as well as quantifiable plasticity in relation to ELA. We first compared mCG and mCAC in distinct genomic features and chromatin states, regardless of clinical grouping. While interactions between DNA methylation and histone marks were described previously in other tissues^63,64^, and in the brain for mCG^63^, here we unravel unforeseen specificities regarding mCAC. First, among the 3 chromatin promoter states (Act-Prom, Wk-Prom, Flk-Prom), mCAC was selectively enriched in Wk-Prom, which was not observed for mCG. Considering that Wk-Prom was relatively depleted in H3K27ac and H3K4me1 compared to the 2 other promoter states, it is possible to hypothesize that these 2 histone modifications may potentially repress mCAC accumulation in brain tissue. A second dissociation consisted in the fact that lower mCAC levels were measured in Str-Trans compared with Wk-Trans regions, while no such difference was observed in the CG context. This may result at least in part from higher levels of H3K36me3 observed in the Str-Trans state. Third, among the 2 tightly compacted chromatin states defined by the repressive mark H3K9me3, PcR and Heteroch, the latter state was characterized by higher DNA methylation in the CG, but not in the CAC, context, as well as by a relative increase in H3K9me3 and a decrease in H3K27me3. While there is currently no data, to our knowledge, supporting a potential interaction between H3K27me3 and non-CG methylation^11^, a role for H3K9me3 can be speculated considering studies of cellular reprogramming. Indeed, *in vitro* dedifferentiation of fibroblasts into induced pluripotent stem cells associates with the restoration of non-CG methylation patterns characteristic of stem cells, except in genomic regions characterized by high levels of H3K9me3^65^. Therefore, the possibility exists that H3K9me3 may be implicated in the regulation of mCAC in the brain, a hypothesis that warrants further investigation.

Finally, we wondered whether mCAC may also contribute to molecular responses to ELA in the brain. We found that similar numbers of differential methylation events could be detected across CAC and CG contexts in ELA subjects, suggesting that both contexts might be sensitive to behavioral regulation. While previous studies already showed that ELA associates with widespread effects on mCG throughout the genome, they were conducted using methodologies primarily designed for the investigation of mCG (methylated DNA immunoprecipitation coupled to microarrays^66^, reduced representation bisulfite sequencing^40^). In comparison, the present WGBS study provides a more comprehensive and unbiased assessment of the overall methylome, and represents, to our knowledge, the first indication in humans that the mCAC form of DNA methylation might be affected by ELA and related depressive phenotypes. This is consistent with recent mouse work^67^ that provided evidence for an effect on non-CG methylation of early-life positive experiences (in the form of environmental enrichment during the adolescence period), suggesting that both beneficial and detrimental experiences may modulate this non-canonical epigenetic mechanism. Importantly, our combined investigation of DNA methylation and histone marks provides further characterization of this form of plasticity. Strikingly, mCAC and mCG changes occurred in genomic regions that appeared distinct at every level of analysis, including genic or methylomic features, individual histone marks, chromatin states, and GO categories. Accordingly, CG-DMRs primarily located among promoter regions and gene bodies, were enriched in H3K4me1, H3K4me3 and H3K27ac, and present across all chromatin states. In comparison, CAC-DMRs were less frequently found in promoters, enriched in H3K36me3 and depleted in H3K9me3, and mostly associated with Quiescent and Wk Trans chromatin states. Furthermore, previous studies on non-CG methylation focused on the comparison of distinct cell types (glial *vs* neuronal cells^12^, or excitatory *vs* inhibitory neurons^24^), and identified parallel and significant differences in both CG and non-CG contexts at common genomic sites. In sharp contrast, regions showing differential methylation as a function of ELA in the present study were clearly context-specific, indicating a more subtle and specific modulation of DNA methylation by ELA than by cell identity. Overall, these results are consistent with a model whereby the cascades of neurobiological adaptations associated with ELA result from and contribute to distinct pathophysiological phenomenon that differentially manifest at the level of CG and CAC sites. It is possible also to speculate that part of these adaptations may result from the impact of ELA on mechanisms that drive the developmental emergence of mCAC. Along this line, a molecular pathway has recently started to be unravelled in the mouse: the methyltransferase Dnmt3a was shown to mediate progressive post-natal accumulation of DNA methylation in the CA context^13^, while *in vivo* recruitment of MeCP2 primarily relies on mCG and mCAC levels (rather than methylation at other contexts, including CAT, CAA, or CAG^28^). These 2 studies suggest that DNMT3a and MeCP2 may be implicated in human in the particular cross-talk that emerges during brain maturation between mCAC and specific histone modifications (H3K27ac, H3K4me1, H3K36me3, H3K9me3). Therefore, future investigations should focus on these marks, and related histone-modifying enzymes, to better decipher the lifelong impact of ELA on molecular epigenetic cross-talks.

In conclusion, the epigenetic and transcriptomic landscape of the lateral amygdala exhibited targeted reconfigurations as a function of ELA. This reprogramming could be detected consistently across multiple epigenetic mechanisms, including the newly recognized form of DNA methylation affecting CAC sites. Future studies will hopefully define the extent to which non-CG methylation at CACs, and potentially at other cytosine contexts, contribute to adaptive and maladaptive encoding of life experiences in the brain.

## Methods

Methods and associated references are available in the online version of the paper.

## Authors contributions

PEL and GT conceived the study. GGC, EM, JY and PEL performed RNA and DNA extractions. TK and TP provided library preparations and next-generation sequencing through the IHEC consortium, and AR prepared ChIP-Seq libraries. MAC, JFT and PEL analyzed histone data. ZA, JFT and PEL analyzed methylation data. JFT and PEL analyzed RNA-Seq data, while the Kallisto analysis was conducted by JCG. AP and JFT conducted the BSmooth differential methylation analysis. MA, LCV, CE, IY, NM and GT provided lab resources and equipment. PEL, MAC and GT prepared the manuscript, and all authors approved its final version.

## Acknowledgments

PEL was supported by fellowships from the ‘Fondation Fyssen’, the Canadian Institutes of Health Research, the American Foundation for Suicide Prevention, the ‘Fondation pour la Recherche Médicale’, and by fundings from the Bettencourt-Schueller Foundation, the Brain Canada Foundation, the ‘Fondation Deniker’, the ‘Congrès Français de Psychiatrie’, the ‘Fondation de France’ (N°Engt:00081244) and the ‘Union Nationale de Familles et Amis de Personnes Malades et/ou Handicapées Psychiques’.

