## Supplementary material for "Non-CG methylation and multiple epigenetic layers associate child abuse with immune and small GTPase dysregulation"

**Supplementary material contents**

1. **Online methods**
2. **Legends of supplementary figures**
3. **Supplementary figures**

#### Online methods

##### Human samples & tissue dissections

Post-mortem lateral amygdala brain tissue was obtained in collaboration with the Quebec Coroner's Office, from the Douglas-Bell Canada Brain bank (douglasbrainbank.ca/, Montreal, Canada). This study included (i) subjects who died suddenly without prolonged agonal state or protracted medical illness, and with no history of psychiatric disorder (Controls, C, N=17), and (ii) subjects with a history of severe child abuse, who died by suicide in the context of a major depressive episode (Early-life adversity, ELA, N=21). Sample characteristics are presented in Supplementary Table1. Groups were matched for age, post-mortem interval (PMI) and brain pH. Psychological autopsies were performed by trained clinicians on both controls and cases, with the informants best-acquainted with the deceased, as described previously[1] and as validated by our group and others[2–7]. Diagnoses were assigned based on DSM IV criteria. Characterization of early-life histories was based on adapted Childhood Experience of Care and Abuse (CECA) interviews assessing experiences of sexual and physical abuse, as well as neglect[8,9], and for which scores from siblings are highly concordant[2,9]. We considered as severe early-life adversity reports of non-random major physical and/or sexual abuse during childhood (up to 15 years). Only cases with the maximum severity ratings of 1 and 2 were included. This information was then complemented with medical charts and coroner records. Ethical approval was obtained from the Institutional Review Board of the Douglas Mental Health University Institute. Written informed consent was obtained from the families of each of the deceased subjects prior to inclusion in the study.

##### Next-Generation Sequencing

WGBS, RNA-Seq and ChIP-seq experiments were carried out by expert technicians at the McGill University and Genome Quebec Innovation Center, following standard operating procedures from the International Human Epigenome Consortium (IHEC, see ihec-epigenomes.org/). Data from the present study will be made publicly available during regular data releases on the consortium data portal.

##### ChIP-seq library preparation

Because of the small size of the amygdala lateral nucleus, and the large amounts of tissue required for multiple immune-precipitations and the ChIP-seq analysis of 6 histone marks, tissue from 17 C and 21 ELA subjects were distributed into 7 ELA and 4 C pools, for an average of 472 mg of tissue available for ChIP-Seq experiments per pool (see Supplementary Table2). Libraries were prepared using the automated protocol for the Kapa HTP Library Preparation Kit (Illumina), and sequencing was performed using the Illumina HiSeq2000, as per the manufacturer's instructions, to achieve at least 30 and 60 million reads for narrow (H3K27ac,

H3K4me3) and broad (H3K27me3, H3K36me3, H3K4me1, H3K9me3) marks, respectively (FigS1a).

**ChIP-seq data processing**

Trimmomatic[10], BWA[11], Picard and deepTools[12] were used to pre-process and align the sequencing reads. Global visualization for the ChIP-seq data was accomplished using IGV[13] and ngs.plot[14]. Inter-sample correlations and hierarchical clustering were achieved using deepTools. Identification of differential enrichment sites for each histone mark was done using diffReps with window size 1000bp, sliding step 100bp and fragment size 200bp[15]. A FDR <10% and p <0.0001 for negative binomial test were used as significance cutoffs. ChromHMM was used to partition the genome into 200bp bins, and annotate them to chromatin states[16]. A 10-state model was chosen and applied to all data sets. A consensus map was first created for general characterization of the amygdala chromatin states, using genomic regions with at least 6/11 samples in accordance for a state. For comparisons of the 2 clinical group, group-specific maps were defined using regions showing at least a 70% agreement between samples (at least 3/4 C pools and 5/7 ELA pools). State transitions (ST) were then defined as regions with differing states between the C and ELA maps. For characterization of the distribution of DS and ST, we used the region_analysis package to annotate them to genomic features[15].

**WGBS library preparation**

Whole-genome sequencing libraries were generated from 700 to 1,000 ng of genomic DNA spiked with 0.1% (w/w) unmethylated λ DNA (Promega) previously fragmented to 300–400 base-pairs (bp) peak sizes using the Covaris focused-ultrasonicator E210. Fragment size was controlled on a Bioanalyzer DNA 1000 Chip (Agilent) and the KAPA High Throughput Library Preparation Kit (KAPA Biosystems) was applied. End repair of the generated dsDNA with 3′- or 5′-overhangs, adenylation of 3′-ends, adaptor ligation and clean-up steps were carried out as per KAPA Biosystems' recommendations. The cleaned-up ligation product was then analysed on a Bioanalyzer High Sensitivity DNA Chip (Agilent) and quantified by PicoGreen (Life Technologies). Samples were then bisulfite converted using the Epitect Fast DNA Bisulfite Kit (Qiagen), according to the manufacturer's protocol. Bisulfite-converted DNA was quantified using OliGreen (Life Technologies) and, based on quantity, amplified by 9–12 cycles of PCR using the Kapa Hifi Uracil+DNA polymerase (KAPA Biosystems), according to the manufacturer's protocol. The amplified libraries were purified using Ampure Beads and validated on Bioanalyzer High Sensitivity DNA Chips, and quantified by PicoGreen. Libraries were run on an Illumina HiSeq 2000 (100bp paired-end), yielding ≈164 million reads/library on average (Fig.S5c), and generating around 6.2 billion reads in total across the whole cohort.

**WGBS data processing**

As previously described[17], in-house generated methylome libraries were aligned using BWA 0.6.1[11] after converting all the reads in bisulfite mode to the human hg19/GRCh37 genome reference. Both reads in a pair were trimmed of any low-quality sequence at their 3' ends (with Phred scale score >=30). Post-process read mappings were made as previously described[17], including clipping 3' ends of overlapping read pairs in both forward and reverse strand mappings, filtering duplicate, low-mapping quality reads, read pairs not mapped at the expected distance based on the library insert size as well as reads with more than 2% mismatches. Methylation calls of individual cytosines in both CG and CAC contexts were extracted using Samtools in mpileup mode. Cytosines overlapping SNPs from dbSNPs (137) and CpGs located within ENCODE DAC blacklisted regions or Duke excluded regions[18] were discarded.

***Methylation data characterization.*** All analyses conducted to characterize the genome-wide abundance and distribution of CG and CAC methylation were done by focusing on cytosines showing a coverage >= 5 (Fig.2 and Fig.S6-9). We used the region_analysis package[15] to assign each cytosine to a genomic feature (Fig.S7), using the Ensembl v75 annotation for consistency with RNA-Sequencing data analysis (see below). MethySeekR was used to call CpG-rich, unmethylated regions (UMR), as well as CpG-poor low-methylated regions (LMR), as described by Burger et al[19].

***Differential methylation analysis.*** Differential methylation analysis was conducted using BSmooth, as described previously[20]. The context of each C was determined, which allowed us to classify each C of the genome as CG or CAC. Methylation levels for each site were estimated by counting the number of reported C ('methylated' reads) divided by the total number of reported C and T ('methylated' plus 'unmethylated' reads) at the same position of the reference genome. To identify differentially methylated regions in the CG context, we performed a strand-independent analysis of CG methylation where counts from the two Cs in a CG and its reverse complement (position *i* on the plus strand and position *i*+1 on the minus strand) were combined and assigned to the position of the C in the plus strand. The summarized methylation estimates of strand-merged CG sites from the 21 ELA and 17 control samples were used to identify differences in methylation, using the R package BSmooth/BSseq[21] at default parameters. To minimize noise in methylation estimates due to low-coverage data, we restricted the differential methylation analysis to CpG sites with coverage of ≥4 sequence reads in at least 10 samples in each condition, which still allowed us to interrogate changes in methylation levels at ~18 million CG and ~39 CAC million sites. The same strategy was applied for differential methylation analysis in the CAC context, except that by definition methylation data originated for each CAC site from one DNA strand only. We identified differentially methylated regions (DMRs) as regions containing at least 5 consecutive CG, or CAC, sites that were significantly differentially methylated using an unpaired Welch t-test ($p < 0.001$) and that exhibited at least a 1% difference in mean methylation levels between ELA and C groups. To rule out potential confounding effects of age and sex (two factors known to contribute to variations in DNA methylation[22,23]), a generalized linear model taking into account these 2 variables was computed on mean methylation levels for each CG- and CAC-DMR. Only those DMRs for which differential methylation between C and ELA subjects remained significant when correcting for age and sex were considered for downstream analyses. Finally, genomic features were attributed to DMRs using the region_analysis package, similar to the annotation of ChIP-Seq DS or ST, while intersections of DMRs with UMRs and LMRs were determined using Bedtools.

**RNA-Sequencing library preparation**

RNA was extracted from homogenized brain samples using the RNeasy Lipid Tissue Mini Kit (Qiagen). Quantity and quality of extracted RNAs were measured using an Agilent 2100 Bioanalyzer. RNA-Sequencing libraries were prepared by expert technicians at the McGill University and Genome Quebec Innovation Center, using IHEC procedures. Briefly, we used the TrueSeq Stranded Total RNA Sample Preparation kit (Illumina), using the Ribo-Zero Gold kit (Illumina) for depletion of ribosomal RNA, followed by first and second strand cDNA synthesis and fragmentation of dsDNA. Then, fragmented DNA was used for A-tailing, adaptor ligation and 12 cycles of PCR amplification. Libraries were quantified using high sensitivity chip on a Labchip (PerkinElmer), quantitative PCR (KAPA Library Quantification, Kapa Biosystems), and PicoGreen (Life Technologies). Three libraries were run per lane of an Illumina HiSeq 2000 (100bp paired-end), yielding ≈54 million reads/library (Fig.S16b).

**RNA-Sequencing data processing**

***Alignment, counting, and differential expression analysis***. As described previously[24,25], we used: FASTX-Toolkit (hannonlab.cshl.edu/fastx_toolkit/links.html) and Trimmomatic[10] for adapter trimming; Bowtie2 for alignment ; TopHat[26] for transcript alignment; HTSeq-count[27] or Kallisto[28] for counting; and DESeq2[29] for differential expression analysis. *Alignment*. Following high-throughput sequencing, 100bp paired-end reads were aligned to the hg19 human genome using TopHat v2.1.0 (tophat.cbcb.umd.edu/) with a mate insert distance of 75 bp (-r) and library type fr-unstranded. Reads passing a mapping quality of at least 50 were used for gene and transcript quantification. *Quantification*. Gene annotations from the Ensembl release 75 were used for gene-level quantification quantification. First, we used HTSeq-count version 0.6.1p1 (www-huber.embl.de/users/anders/HTSeq/doc/overview.html), using the intersection-nonempty mode, and results were combined to form a count matrix of 20,893 transcribed

RNAs across 50 samples. As an alternative strategy to HTSeq-count, we also processed reads through the pseudo-aligner Kallisto. Here, expression counts were obtained for isoforms using Kallisto (v. 0.43.0). Then, the tximport (v. 1.0.3) R package was used to reconstruct gene-level counts using the isoform-level counts generated by Kallisto. *Differential expression analysis*. Genes with no mapped fragments were removed from the analysis. Furthermore, genes with low counts were removed by keeping only those with at least 20 counts per subject in average. Using HTSeq-count, differential expression analysis was performed using the DESeq2 general linear model (GLM) using the following covariates: gender[30], age[31], pH, PMI, and RIN[32], based on previous literature documenting their impact on human brain RNA-Seq datasets.

***Gene Set Enrichment Analysis (GSEA).*** GSEA was performed as previously described[25,33]. $Log_2$ fold changes were obtained for each gene from the differential gene expression analysis. Genes were ranked based on their fold changes where genes with the highest positive fold changes were at the top of the list and those with the lowest negative fold changes were at the bottom of the list. The ranked gene list was then used as an input for the GSEAPreranked tool, with the "classic" enrichment score calculation option selected. The C2 curated gene sets molecular signatures database was used to identify enriched gene sets.

**Deconvolution of cellular composition**

To assess the abundance of various cell types in our amygdala samples, we used BSEQ-sc[34] and the CIBERSORT[35] algorithm, and applied these to both our RNA-Sequencing and WGBS data. For deconvolution of these data, we used as reference 'signatures': i) for gene expression: a matrix built from single-nuclear RNA-Sequencing data recently generated by our group using prefrontal cortical tissue[36] (archived on GEO Datasets under the reference series GSE144136), and analysed using unsupervised graph-based clustering[37], and ii) for DNA methylation: the non-CG methylation matrix generated by Luo and colleagues[38] using single-nucleus methyl-cytosine sequencing. Relative fractions of cells were computed and are displayed in FigS14.

**Gene ontology**

We used GREAT v3.0.0[39] to identify the enrichment of gene categories in differential sites (DS) or state transition sites (ST) obtained from ChIP-Seq experiments, and for DMRs from WGBS experiments. DS, ST and DMRs were associated with genes using the default proximal (5kb upstream, 1kb downstream of TSS) and distal (+/- 1Mb of TSS) definition of regulatory regions. Biological process and molecular function gene categories were kept if they passed both the hypergeometric and binomial tests with a fold enrichment ≥1.5 and FDR Q≤0.1. Significant GO terms with less than 5 genes associated with ST, DS or DMRs were discarded. To account for the recurrence of terms across multiple combinations of ST, we calculated a co-occurrence score for each GO term, consisting of the sum of the –log10 of the binomial p-value for each ST enriched in this term, as described by Feng et al[40].

**Supplementary Figures legends.**

**Supplementary Figure 1. Quality controls for chromatin immunoprecipitation sequencing (ChIP-Seq). (a)** ChIP-Seq libraries were prepared and sequenced by expert technicians at the Genome Québec Innovation Center, in the framework of the International Human Epigenomics Consortium (IHEC). We compared psychiatrically healthy controls (C) and subjects with a history of early-life adversity (ELA), and analyzed 4 'broad' (H3K9me3, H3K27me3, H3K36me3, H3K4me1) and 2 'narrow' (H3K4me3, H3K27ac) marks, for which we aimed at sequencing roughly 60 and 30 million reads per library, respectively, as per the IHEC consortium's guidelines. A two-way ANOVA indicated that there was no difference among C and ELA groups in terms of sequencing depths [$F(1,63)=0.52$; $p=0.47$]. Values are mean±sem. **(b)** Quality controls analyses showed that samples for narrow marks showed greater than 0.8 and 1.05 Relative Strand Cross and Normalized Strand Cross correlations, respectively, thereby meeting ENCODE consensus thresholds for quality control[41].

**Supplementary Figure 2. Comparison of amygdalar histone profiles for individual marks with datasets from inferior temporal lobe (Inf Temp), anterior caudate (Ant Caud), and peripheral blood mononuclear cells (Blood).** Data were downloaded from the Roadmap Epigenomics Consortium (ncbi.nlm.nih.gov/geo/roadmap/epigenomics/) and compared to each C and ELA amygdalar sample from the present study, using deepTools Pearson correlations and unsupervised clustering[42], for each of the 6 histone marks: **(a)** H3K4me1; **(b)** H3K27ac; **(c)** H3K4me3; **(d)** H3K36me3; **(e)** H3K9me3; **(f)** H3K27me3. Accession numbers: 1) for inferior temporal lobe: datasets GSM772995 (H3K27ac), GSM772993 (H3K27me3), GSM772982 (H3K36me3), GSM772992 (H3K4me1), GSM772996 (H3K4me3), GSM772994 (H3K9me3); 2) for anterior caudate: datasets GSM772832 (H3K27ac), GSM772827 (H3K27me3), GSM772828 (H3K36me3), GSM772830 (H3K4me1), GSM772829 (H3K4me3), GSM772831 (H3K9me3); 3) for blood mononuclear cells: datasets GSM1127145 (H3K27ac), GSM1127130 (H3K27me3), GSM1127131 (H3K36me3), GSM1127143 (H3K4me1), GSM1127126 (H3K4me3), GSM1127133 (H3K9me3). Of note, H3K4me1, H3K27ac and H3K27me3 better discriminated between tissue types than the 3 other marks.

**Supplementary Figure 3. Characterization of amygdalar chromatin states defined using ChromHMM, and comparison with external datasets from human brain hippocampus and gastric tissues. (a)** A snapshot of the ChromHMM 10-state model that was generated using amygdalar data for 6 histone marks is shown. **(b)** The graph depicts the number of genomic regions identified for each chromatin state; each region is defined as continuous genomic bins showing a similar combination of individual histone marks in the consensus ChromHMM model (see Methods). **(c)** Distribution of genomic size of regions identified for each chromatin state. **(d)** Methylation levels in the CAC context (y-axis) were plotted against the region size (x-axis) for the 3 Promoter states: Active Promoter (Act Prom, upper panel), Weak Promoter (Wk Prom, middle panel), and Flanking Promoter (Flk Prom, lower panel). **(e-f)** Comparison of the amygdalar ChromHMM 10-state consensus model generated here using amygdalar tissue with similar chromHMM models generated by the NIH Roadmap. Brain hippocampus and gastric tissue datasets were downloaded from the Roadmap Epigenomics Project portal (Epigenome ID E071 and E094 respectively, for the core 15-state model, egg2.wustl.edu/roadmap/web_portal/chr_state_learning.html) and compared using Bedtool's Jaccard coefficient to our consensus map (generated using ChromHMM, see *Methods*). Note that our amygdala chromatin model more strongly correlates with that of the brain hippocampus than with gastric tissue, as expected.

**Supplementary Figure 4. Relationship between chromHMM chromatin state and gene expression levels.** For each gene (+/- 3kb), the percentage of its genomic span covered by each chromatin state was computed. Values were then ordered by gene expression level and averaged over bins of 500 genes (values depicted are mean ± sem). Here are shown the average percentage coverage of genes by **(a)** Weak transcription (Wk Trans) and Strong transcription (Str Trans), **(b)** Genic enhancer (Gen Enh) and Enhancer (Enh), **(c)** Active promoter (Act Prom), Weak promoter (Wk Prom) and Flanking promoter (Flk Prom) states. These chromatin states exhibited expected correlations with gene expression: (i) the Wk Trans state was more frequently observed in gene bodies of lowly expressed genes compared with the Str Trans state, (ii) enhancer states were enriched in genes with strong expression and (iii) the Wk Prom state was more frequently observed in lowly expressed genes compared with the Act Prom state.

**Supplementary Figure 5. Quality controls for whole-genome bisulfite sequencing libraries (WGBS).** To compare DNA methylation patterns among psychiatrically healthy individuals (controls, C) and subjects with a history of severe child abuse (early-life adversity, ELA), WGBS libraries were prepared. **(a)** Bisulfite conversion efficiencies were measured for each DNA sample using spiked-in unmethylated lambda DNA, and were similar between groups: C: 99.3±0.07%, ELA: 99.2±0.06% ($t=0.63$, $p=0.53$). **(b)** In addition, over-conversion (i.e. methylated cytosines converted to uraciles during bisulfite conversion) was determined experimentally using spiked-in fully methylated pUC19 DNA, and similar values were observed between groups: C: 5.6±0.10%, ELA: 5.7±0.05% ($t=0.19$, $p=0.85$). **(c)** Each WGBS library was then sequenced on 1 lane of a HiSeq 2000 (100 base pair, paired-end sequencing) at the Génome Québec Innovation Center, yielding similar sequencing depth across groups: C: 164±3 million reads, ELA: 163±3 million reads ($t=0.15$, $p=0.88$). **(d)** During the processing of raw sequencing data, duplicates were removed from downstream analysis, and results indicated similar diversity (t=1.18, p=0.25) among libraries from the C (C:22.5±1.2%) and ELA (C:20.8±0.9%) groups. **(e)** The graphs depicts the number of CG sites that met distinct average coverages among the 38 WGBS libraries. For the characterization of genome-wide abundance and distribution of CG and CAC methylation levels (Fig.2 and Fig.S6-9), we focused on cytosines with a coverage >= 5. Abbreviations: cov., coverage. Values are mean±sem.

**Supplementary Figure 6. Distribution of non-CG DNA methylation in the human brain lateral amygdala. (a)** The graph, taken from Mo et al[43], depicts the distribution in the mouse of non-CG methylation, measured in 3 neuronal subtypes: glutamatergic neurons (Exc), and parvalbumin-expressing (PV) and vasoactive intestinal peptide (VIP)-expressing inhibitory neurons. **(b)** A very similar distribution of genome-wide average non-CG methylation levels was observed in the human brain lateral amygdala, in healthy controls (C) and subjects with a history of child abuse (early-life adversity, ELA). While most cytosines in non-CG contexts were unmethylated, a minority of these positions nevertheless showed methylation levels between 5 and 25% **(c)**, whatever the 3-letter context considered, with a peak between 15 to 20%. These results indicate that while different numbers of cytosines might be methylated across various non-CG contexts, the abundance of DNA methylation (i.e., the proportion of cells affected) at those sites seems relatively homogeneous. Values are mean±sem.

**Supplementary Figure 7. Distribution of CG and CAC DNA methylation among distinct genomic features and chromosomes. (a)** CAC and CG methylation levels were computed across distinct genomic features defined using the region_analysis package[15]. In the CG context, DNA methylation levels strongly varied as a function of the genomic feature [F(7,252)=953; p<0.0001], while there was no significant difference [F(1,36)=0.50; p=0.48] among psychiatrically healthy individuals (C) and subjects with a history of early-life adversity (ELA). Post-hoc comparisons confirmed that, as expected, lowest CG methylation levels were observed in promoter regions, in particular within a 250-base pair distance from the TSS (ProximalPromoter), where methylation levels were significantly lower than in any other gene feature (p<0.0001). In the CAC context, similar significant and non-significant effects were found for genomic features [F(7,252)=917; p<0.0001] and for clinical grouping [F(1,36)=0.03; p=0.85], respectively. In contrast with the CG context, lowest CAC methylation levels were observed in pericentromeric regions (defined by region_analysis as regions located between the boundary of a centromere and the closest gene minus 10kbp of that gene's regulatory region[15]), which showed strongly significant differences with all other features (p<0.0001). Values are mean±sem. **(b)** CAC and CG methylation levels were computed in each chromosome across the whole cohort. As expected, DNA methylation levels were much higher in the CG than in the CAC context (2-way ANOVA; context effect: [F(1,1850)=1238951; p<0.0001]), and methylation abundance strongly varied among chromosomes in each context (chromosome effect: [F(24,1850)=2673; p<0.0001]). As expected also, methylation was extremely low in the mitochondrial genome in both the CG and CAC contexts. Values are mean±sem.

**Supplementary Figure 8. Correlations between the expression of genes and DNA methylation levels in their sense or antisense strands, in the CG and CAC contexts.** The 1000 most highly (top1000) and 1000 most lowly expressed genes were identified using RNA-Sequencing data, and compared for abundance of DNA methylation in: **(a,d)** both DNA strands; **(b,e)** the strand where genes are located (sense strand), and **(c,f)** the strand antisense to the one where genes are located. Negative correlations between DNA methylation and gene expression were observed in all cases: **(a)** [F(1,74)=736.1; p<2E-16] **(b)** [F(1,74)=742.8; p<2E-16] **(c)** [F(1,74)=615.1; p<2E-16] **(d)** [F(1,74)=145.3; p<2E-16] **(e)** [F(1,74)=119.2; p<2E-16] **(f)** [F(1,74)=142.6; p<2E-16] (2-way repeated measures ANOVA, main effects of gene category, top1000 versus bottom1000 averaged over 100 bins). Results therefore indicate that gene expression is predicted to the same extent by mCAC on either strand, at least for the coverage achieved in this study.

**Supplementary Figure 9. Annotation of DNA methylation features in the human brain lateral amygdala. (a)** Unmethylated (UMR) and lowly methylated (LMR) genomic regions were identified using the methylseekR algorithm, as described in[19]. No partially methylated domains (PMDs) were identified, similar to previous studies on mouse retina[44], human embryonic stem cells and neural progenitor cells[45]. **(b-c)** LMR (n=115254) and UMR (n=21267) showed methylation levels in the CG context (b) and size (c) consistent with previous reports[19], while UMR (mean ± sem = 2516 ± 41 bp) were generally larger than LMR (738 ± 107 bp). Dashed and dotted lines represent medians and quartiles, respectively. **(d)** Among all CpG islands in the human genome (RefSeq reference, n= 28691), a vast majority (n=19193; n=66.9%) intersected with CpG-dense UMR**,** while a small minority corresponded to LMR (n=2743; 9.6%), as described previously for other tissue types[45]. **(e)** Methylation in the CG (mCG) and CAC (mCAC) contexts were computed across RefSeq genomic features, UMR and LMR, and found to strongly differ, as expected. While mCAC levels are much lower than mCG levels, a similar pattern of variation is observed across the 2 cytosine contexts. **(f)** mCG (left panel) and mCAC (right panel) levels were computed in each intersection among RefSeq genomic features, LMR, or UMRn and chromatin states (identified using ChromHMM, see *Methods*), across our entire cohort. Of note, while mCG abundance in LMRs remained constant between 25 and 35% across all chromatin states, mCAC was more variable, and notably lower in Polycomb Repressed (PcR), heterochromatin (Heterochr) and Strong Transcription (Str Trans) states. Also, mCAC was more abundant in Wk Trans than in Str Trans within genes, consistent with the fact that Wk Trans was more abundant in lowly expressed genes (see FigS9a), and previous reports[46].

**Supplementary Figure 10. Histone profiles at lowly methylated regions (LMR) and unmethylated regions (UMR). (a)** Enrichment of chromatin states at and around positions of UMR (left panel) and LMR (right panel) were evaluated using chromHMM's NeighborhoodEnrichment function. As expected, UMRs co-located with Weak and Active Promoters (Wk Prom, Act Prom), and were flanked by the Flanking Promoter (Flk Prom) state. In contrast, LMRs were consistently surrounded by Str Enh and Enh states, which spanned 2 and 1 Kb around LMR positions, respectively, consistent with their role as distinct regulatory element[45]. **(b-g)** Average enrichments of ChIP-seq reads over input are shown for each histone mark within LMR and UMR, as well as in flanking up- and down-stream regions (+/- 1 Mb). Patterns are consistent with those reported by Stadler et al[45] for 2 marks (H3K4me1, H3K4me3), and provide new information related to the other 4 marks: i) LMR and UMR associated with increased levels of H3K4me1, H3K4me3 and H3K27ac, and these effects were more pronounced for UMR than for LMR, consistent with the fact that CpG islands showed a stronger overlap with UMR than with LMR (Fig.S7d); ii) The other 3 marks (H3K36me3, H3K9me3, H3K27me3) were depleted within UMR and LMR; iii) interestingly, while LMR regions showed uniform enrichment or depletion throughout their whole genomic span, more complex patterns were observed for UMR. Their enrichment for H3K4me1 showed a biphasic pattern, and preferentially affected their 5' and 3' shores, very similar to results obtained by Stadler et al in mouse embryonic stem cells; further, this biphasic enrichment was also present for H3K4me3 and H3K27ac, albeit to a lesser extent, and was mirrored by a biphasic H3K9me3 depletion in UMR shores.

**Supplementary Figure 11. Distributions of histone reads across gene bodies and differential sites (DS).** For each histone mark, the figure depicts the distribution of reads (average enrichment of ChIP-seq reads over input) across all gene bodies (All Genes). In addition, and as a control, we also analyzed their distribution at genomic sites where Up- and Down-DS were identified between control (C) and early-life adversity (ELA) groups. Values are mean±sem.

**Supplementary Figure 12. Identification of genomic features where histone differential sites (DS) were localized. (a)** Localization of DS (identified using diffRep, see *Methods*) among distinct genomic features (defined using region_analysis[15]), for each histone mark. The observed distributions were significantly different across histone marks (df=25, $\chi^2$ =1244, p<0.001). In addition, comparisons between observed and expected (genome-wide distribution of reads among genomic features in the 2 C and ELA groups combined) distributions showed that, for each type of histone modification, ELA-associated DS were non-randomly located in specific genomic features: H3K27ac (df=1, $\chi^2$ =81.2, p<0.001); H3K27me3 (df=1, $\chi^2$ =42.3, p<0.001); H3K36me3 (df=1, $\chi^2$ =138, p<0.001); H3K4me1 (df=1, $\chi^2$ =287, p<0.001); H3K4me3 (df=1, $\chi^2$ =90.9, p<0.001); H3K9me3 (df=1, $\chi^2$ =22.6, p<0.001). **(b)** Analysis of the directionality of DS showed that ELA more frequently associated with decreases (Down-DS) than increases (Up-DS) in read density, as found for 4 marks: H3K4me1 (df=1, $\chi^2$ =231, p<0.001); H3K4me3 (df=1, $\chi^2$ =73, p<0.001); H3K36me3 (df=1, $\chi^2$ =345, p<0.001); H3K27me3 (df=1, $\chi^2$ =228, p<0.001). DS were equally distributed among Up- and Down-DS for the 2 remaining marks: H3K27ac (df=1, $\chi^2$ =0.13, p=0.19); H3K9me3 (df=1, $\chi^2$ =1.7, p=0.19). **(c)** Integrin signaling is enriched across multiple histone changes and state transitions. GREAT pathway analysis using MsigDB showed recurrent enrichment of the integrin signaling pathway across six types of state transitions, as well as for H3K27ac down-DS (differential sites). Each analysis passed hypergeometric and binomial testing (fold change ≥ 1.5 and Q ≤ 0.1 for both tests). Negative logarithmic P-value is shown for the binomial test**.** Chromatin states: Str-Trans, strong transcription; Wk-Trans, weak transcription; Str-Enh, strong enhancer; Enh, enhancer.

**Supplementary Figure 13. Comparison of main metrics for CG and CAC differentially methylated regions (DMR).** DMRs were identified using BSmooth (see *Methods*). **(a)** Compared to CG-DMRs, CAC-DMRs were composed of slightly fewer cytosines (CG: 7.63±0.11; CAC: 7.14±0.08; [t(1,1614.1)=3.63, p=2.9E-04]), and **(b)** were smaller (CG: 321±7 bp; CAC: 245±4 bp; [t(1,1494.9)=9.40, p<2.2E-16]). The amplitude of methylation changes observed in subjects from the ELA group was smaller in the CAC than in the CG context, as shown by **(c)** smaller % changes in methylation levels (CG: 7.75±0.05%; CAC: 4.6±0.03%; [t(1,1338.4)=58.89], p<2.2E-16), and **(d)** smaller areaStat values (a metric measuring the statistical strength of methylation changes among cytosines composing each DMR[21]; CG: 32.3±0.5%; CAC: 28.9±0.4%; [t(1,1575.1)=5.45], p=5.8E-08). Values are mean±sem. **(e)** Visualization of all CG- and CAC-DMRs across the human genome using chromPlot. There was absolutely no intersection between the 2 types of DMRs.

**Supplementary Figure 14. Deconvolution of WGBS and RNA-Sequencing data. (a)** Consistent with the cortical nature of the lateral nucleus of the amygdala and with estimates from neuroanatomical studies[47,48], RNA-Seq deconvolution using BSEQ-sc Cibersort[35] indicated that the neuronal population in our samples was composed of 80% excitatory and 20% inhibitory neurons. Importantly, we found no differences in cell-type composition between

C and ELA groups (2-way ANOVA; cell-type effect $p<0.0001$; group effect, $p=0.61$), whether at the level of 5 major cell types (excitatory and inhibitory neurons, microglia, astrocytes, oligodendroglia, endothelial cells) **(b),** or when considering the full spectrum of 26 cell-type clusters that were identified **(c)**. We then deconvoluted WGBS data using the same Cibersort algorithm and, to our knowledge, the only available dataset for single-cell methylomes in the human brain[38]. Despite the fact that only non-CG methylation levels, but not CG methylation, are available from the later study, we nevertheless observed a significant correlation between the estimated proportion of excitatory neurons in this second approach, and the proportion of the most abundant population of excitatory neurons (Ex_5_L5) observed during deconvolution of RNA-Seq data **(d)**, indicating some convergence between the 2 deconvolutions. **(e)** Importantly, consistent with RNA-Sequencing data, no significant difference in cell-type proportions were identified among C and ELA groups in WGBS deconvolution (2-way ANOVA; cell-type effect $p<0.0001$; group effect, $p=0.11$).

**Supplementary Figure 15. Comparison of histone mark profiles at CG and CAC DMRs. (a-f)** Average enrichments of ChIP-seq reads over input are shown for each histone mark around each type of DMRs (CG, green; CAC, orange): (a) H3K4me1, (b) H3K9me3, (c) H3K4me3, (d) H3K36me3, (e) H3K27ac, and (f) H3K27me3. Histone reads enrichment significantly varied within DMRs and flanking regions (+/- 2 kilobases, kb) for the 6 marks (2-way repeated measures ANOVA, main effects of cytosine position, x-axis; $p<0.0001$). Also, significant differences in histone reads density (y-axis) among the 2 cytosine contexts were observed for 4 marks (H3K27ac, H3K4me3, H3K27me3, H3K4me1; $p<0.05$), but not for H3K9me3 ($p=0.08$) or H3K36me3 ($p=0.57$). Importantly, significant interactions between histone reads density and cytosine position within DMRs were observed for all marks ($p<0.0001$). In particular, post-hoc comparisons confirmed that, compared with their flanking regions, CAC-DMRs were significantly enriched for H3K36me3 and depleted in H3K9me3; in contrast, CG-DMRs were characterized by significant enrichments for H3K4me1, H3K4me3, and H3K27ac, and a depletion in H3K36me3 ($p<0.0001$ for each post-hoc comparison). Values are mean±sem. **(g)** Enrichment of each chromatin state over CG and CAC differentially methylated regions (DMRs). The figure depicts the enrichment of each chromatin state (as identified using ChromHMM and the combination of all 6 histone marks, see main text) among genomic regions corresponding to CG- and CAC-DMRs, compared to their relative abundance in the overall human genome. CG-DMRs shown in blue, CAC-DMRs shown in red. Chromatin states: Act-Prom, active promoter; Wk-Prom, weak promoter; Flk-Prom, flanking promoter; Str-Trans, strong transcription; Wk-Trans, weak transcription; Str-Enh, strong enhancer; Enh, enhancer; PcR, polycomb repressed; Hetero, heterochromatin.

**Supplementary Figure 16. RNA-Sequencing quality controls.** Total RNAs extracted from lateral amygdala tissue (controls, C, n=17; early-life adversity, ELA, n=21) were used for the preparation of RNA-Sequencing libraries and processed in parallel. **(a)** RNA integrity values (RIN) were not significantly different across RNA samples extracted from C (n=17) and ELA (n=21) subjects (Mann-Whitney U=171; p=0.83). **(b)** The number of reads sequenced in each library was similar across C and ELA groups (t-test t=0.72; p=0.48). **(c-d)** Similarly, there was no significant difference in percentages of duplicates (t=1.25; p=0.22) nor in alignment rate (t=0.51; p=0.62) between the 2 groups. Values are mean±sem.

**Supplementary Figure 17. RNA-Sequencing results**. **(a)** Very similar results (r=0.82, t=199.69, p< 2.2E-16) were obtained using 2 distinct bio-informatic pipelines for the analysis of RNA-Sequencing data. Raw reads were aligned & counted using either HTSeq-count or Kallisto, followed by the analysis of differential expression between C and ELA groups using DESeq2. **(b)** Gene Set Enrichment Analysis (GSEA) of RNA-Sequencing data. Genes were ranked based on their $\log_2$ fold changes ('Ranked list metric', from the differential gene expression analysis between C and ELA groups, represented in grey in the lower portion of each panel); genes with highest positive fold changes (in red, upregulated in the ELA group) were at the extreme left of the distribution, and those with lowest negative fold changes (in blue, downregulated in the ELA group) were at the extreme right of the distribution. A running enrichment score (green line, upper portion of each panel) was computed for gene sets from the C2 MSigDB curated molecular signatures database, and used to identify enriched gene sets[33]. Depicted are the 2 single gene sets that achieved the highest normalized enrichment scores (with the middle portion of each panel showing vertical black lines where members of the gene set appear in the ranked list of genes). As shown in the left panel, a collection of genes related to oligodendrocytes and myelin physiology, which we recently found downregulated in the anterior cingulate cortex of subjects with a history of ELA[25], showed an opposite upregulation in the lateral amygdala (normalized enrichment score=2.91; FWER q-value<0.05), suggesting that opposed transcriptional adaptations might occur as a function of ELA between cortical and subcortical structures in this glial population. This finding was reinforced by the second-best gene list (right panel), which was significantly enriched for upregulated genes in our amygdala data (normalized enrichment score=2.75; FWER q-value<0.05). The later gene collection was previously associated with depression in the middle temporal gyrus[49], and was found enriched in myelin-related genes, suggesting that similar stress-related transcriptomic changes may affect oligodendrocytes among distinct portions of the temporal lobe. **(c)** Identification of Gene ontology (GO) processes most consistently affected by early-life adversity (ELA), as identified by the combined analysis of individual histone marks, chromatin states, DNA methylation, and gene expression (see main text).

**FigureS1**

**a**

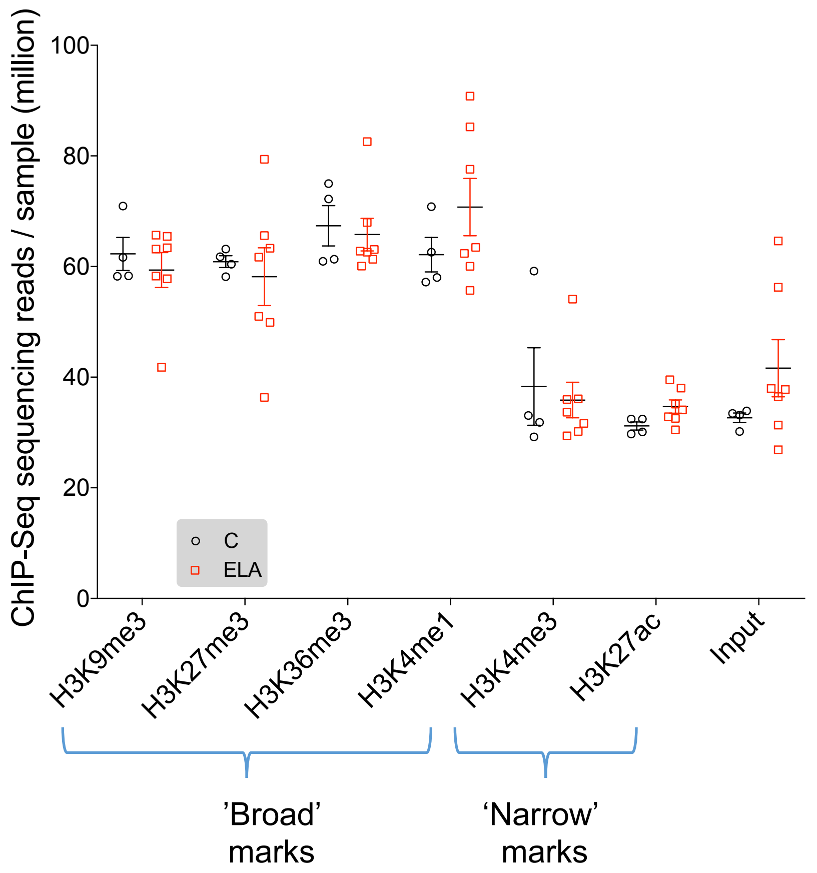

ChIP-Seq sequencing reads / sample (million)
100
80
60
40
20
0
C
ELA
H3K9me3
H3K27me3
H3K36me3
H3K4me1
H3K4me3
H3K27ac
Input
'Broad' marks
'Narrow' marks


**b**

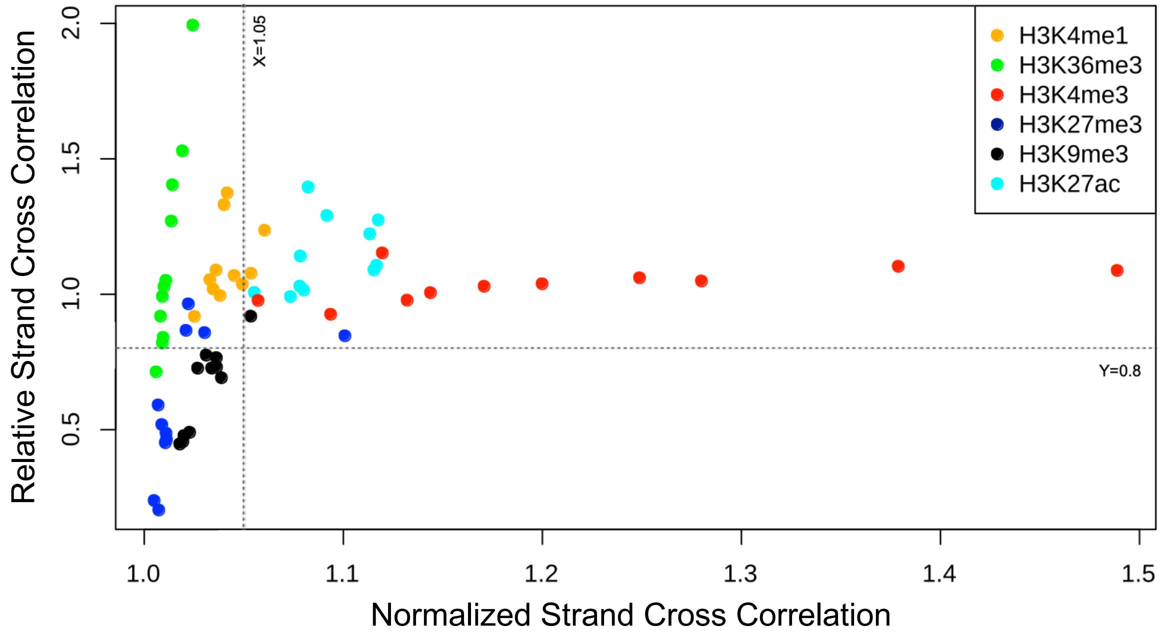

Relative Strand Cross Correlation
2.0
1.5
1.0
0.5
X=1.05
Y=0.8
H3K4me1
H3K36me3
H3K4me3
H3K27me3
H3K9me3
H3K27ac
1.0
1.1
1.2
1.3
1.4
1.5
Normalized Strand Cross Correlation

FigureS2

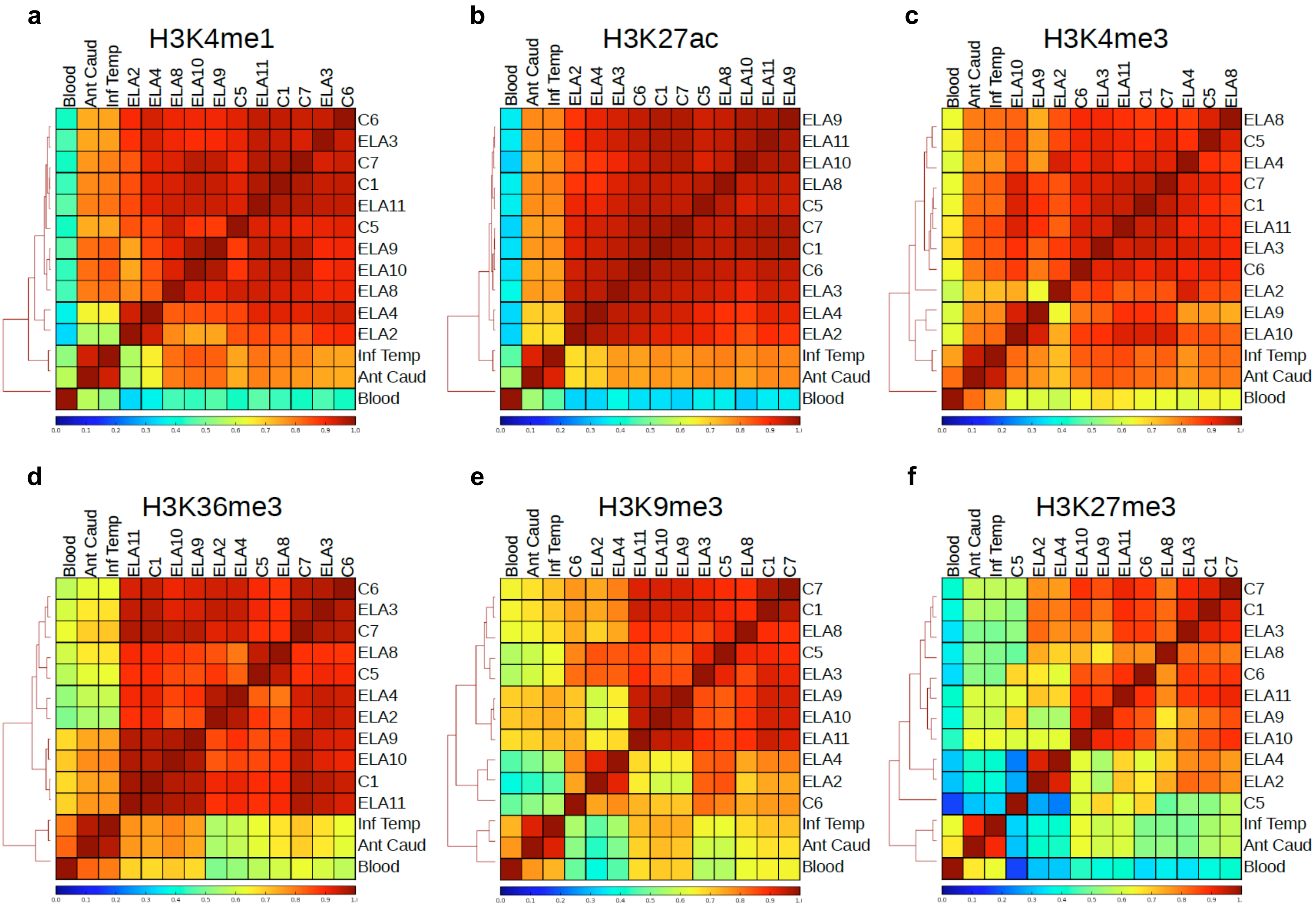
a
H3K4me1
Blood
Ant Caud
Inf Temp
ELA2
ELA4
ELA8
ELA10
ELA9
C5
ELA11
C1
C7
ELA3
C6
b
H3K27ac
Blood
Ant Caud
Inf Temp
ELA2
ELA4
ELA3
C6
C1
C7
C5
ELA8
ELA10
ELA11
ELA9
c
H3K4me3
Blood
Ant Caud
Inf Temp
ELA10
ELA9
ELA2
C6
ELA3
ELA11
C1
C7
ELA4
C5
ELA8
d
H3K36me3
Blood
Ant Caud
Inf Temp
ELA11
C1
ELA10
ELA9
ELA2
ELA4
C5
ELA8
C7
ELA3
C6
e
H3K9me3
Blood
Ant Caud
Inf Temp
C6
ELA2
ELA4
ELA11
ELA10
ELA9
ELA3
C5
ELA8
C1
C7
f
H3K27me3
Blood
Ant Caud
Inf Temp
C5
ELA2
ELA4
ELA10
ELA9
ELA11
C6
ELA8
ELA3
C1
C7

### FigureS3

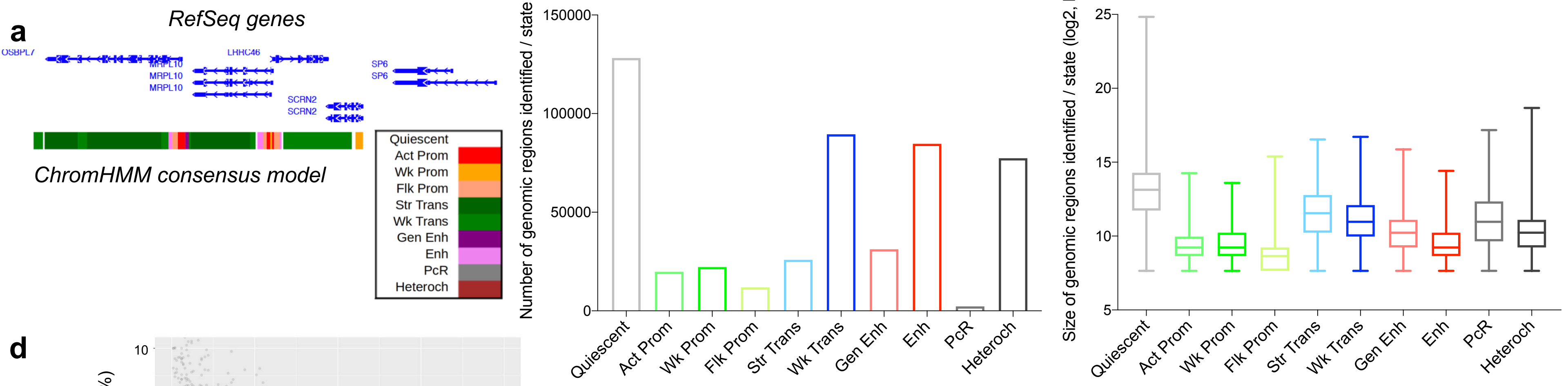


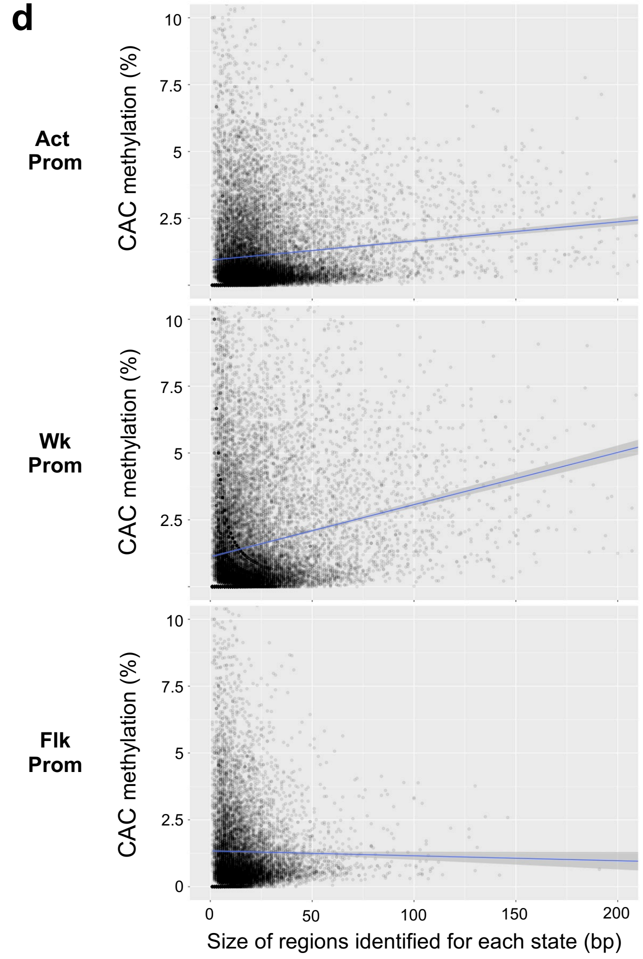


**e** Brain amygdala chromatin states

Brain hippocampus chromatin states

| | Act Prom | Wk Prom | Flk Prom | Str Trans | Wk Trans | Gen Enh | Enh | Heteroch | PcR | Quiescent |
|---|---|---|---|---|---|---|---|---|---|---|
| 1_TssA | 0.29 | 0.32 | 0.02 | 0.00 | 0.00 | 0.00 | 0.00 | 0.00 | 0.00 | 0.00 |
| 2_TssAFlnk | 0.18 | 0.04 | 0.14 | 0.00 | 0.00 | 0.05 | 0.02 | 0.00 | 0.00 | 0.00 |
| 3_TxFlnk | 0.01 | 0.00 | 0.03 | 0.00 | 0.00 | 0.00 | 0.00 | 0.00 | 0.00 | 0.00 |
| 4_Tx | 0.00 | 0.00 | 0.00 | 0.44 | 0.04 | 0.01 | 0.00 | 0.00 | 0.00 | 0.00 |
| 5_TxWk | 0.00 | 0.00 | 0.00 | 0.15 | 0.37 | 0.03 | 0.05 | 0.01 | 0.00 | 0.02 |
| 6_EnhG | 0.00 | 0.00 | 0.02 | 0.01 | 0.00 | 0.07 | 0.01 | 0.00 | 0.00 | 0.00 |
| 7_Enh | 0.01 | 0.00 | 0.01 | 0.00 | 0.02 | 0.26 | 0.26 | 0.00 | 0.00 | 0.00 |
| 8_ZNF/Rpts | 0.00 | 0.00 | 0.00 | 0.00 | 0.00 | 0.00 | 0.00 | 0.03 | 0.02 | 0.00 |
| 9_Het | 0.00 | 0.00 | 0.00 | 0.00 | 0.00 | 0.00 | 0.00 | 0.26 | 0.02 | 0.01 |
| 10_TssBiv | 0.01 | 0.06 | 0.00 | 0.00 | 0.00 | 0.00 | 0.00 | 0.00 | 0.02 | 0.00 |
| 11_BivFlnk | 0.00 | 0.03 | 0.01 | 0.00 | 0.00 | 0.00 | 0.00 | 0.00 | 0.02 | 0.00 |
| 12_EnhBiv | 0.00 | 0.00 | 0.00 | 0.00 | 0.00 | 0.00 | 0.00 | 0.00 | 0.03 | 0.00 |
| 13_ReprPC | 0.00 | 0.01 | 0.00 | 0.00 | 0.00 | 0.00 | 0.00 | 0.00 | 0.11 | 0.01 |
| 14_ReprPCWk | 0.00 | 0.00 | 0.00 | 0.00 | 0.00 | 0.00 | 0.00 | 0.01 | 0.01 | 0.10 |
| 15_Quies | 0.00 | 0.00 | 0.00 | 0.00 | 0.04 | 0.00 | 0.01 | 0.04 | 0.00 | 0.77 |

**f**

Gastric tissue chromatin states

| | Act Prom | Wk Prom | Flk Prom | Str Trans | Wk Trans | Gen Enh | Enh | Heteroch | PcR | Quiescent |
|---|---|---|---|---|---|---|---|---|---|---|
| 1_TssA | 0.21 | 0.29 | 0.03 | 0.00 | 0.00 | 0.00 | 0.00 | 0.00 | 0.00 | 0.00 |
| 2_TssAFlnk | 0.02 | 0.04 | 0.05 | 0.00 | 0.00 | 0.00 | 0.00 | 0.00 | 0.01 | 0.00 |
| 3_TxFlnk | 0.00 | 0.00 | 0.01 | 0.00 | 0.00 | 0.00 | 0.00 | 0.00 | 0.00 | 0.00 |
| 4_Tx | 0.00 | 0.00 | 0.00 | 0.24 | 0.04 | 0.02 | 0.01 | 0.00 | 0.00 | 0.00 |
| 5_TxWk | 0.01 | 0.00 | 0.01 | 0.17 | 0.25 | 0.05 | 0.06 | 0.01 | 0.00 | 0.06 |
| 6_EnhG | 0.00 | 0.00 | 0.01 | 0.01 | 0.01 | 0.02 | 0.01 | 0.00 | 0.00 | 0.00 |
| 7_Enh | 0.02 | 0.02 | 0.02 | 0.00 | 0.02 | 0.04 | 0.07 | 0.00 | 0.01 | 0.01 |
| 8_ZNF/Rpts | 0.00 | 0.00 | 0.00 | 0.00 | 0.00 | 0.00 | 0.00 | 0.02 | 0.03 | 0.00 |
| 9_Het | 0.00 | 0.00 | 0.00 | 0.00 | 0.00 | 0.00 | 0.00 | 0.09 | 0.04 | 0.01 |
| 10_TssBiv | 0.00 | 0.00 | 0.00 | 0.00 | 0.00 | 0.00 | 0.00 | 0.00 | 0.00 | 0.00 |
| 11_BivFlnk | 0.00 | 0.00 | 0.00 | 0.00 | 0.00 | 0.00 | 0.00 | 0.00 | 0.01 | 0.00 |
| 12_EnhBiv | 0.00 | 0.00 | 0.00 | 0.00 | 0.00 | 0.00 | 0.00 | 0.00 | 0.00 | 0.00 |
| 13_ReprPC | 0.00 | 0.00 | 0.00 | 0.00 | 0.00 | 0.00 | 0.00 | 0.00 | 0.00 | 0.00 |
| 14_ReprPCWk | 0.00 | 0.01 | 0.00 | 0.00 | 0.00 | 0.00 | 0.00 | 0.00 | 0.04 | 0.01 |
| 15_Quies | 0.00 | 0.00 | 0.00 | 0.00 | 0.05 | 0.01 | 0.02 | 0.05 | 0.00 | 0.77 |

### FigureS4

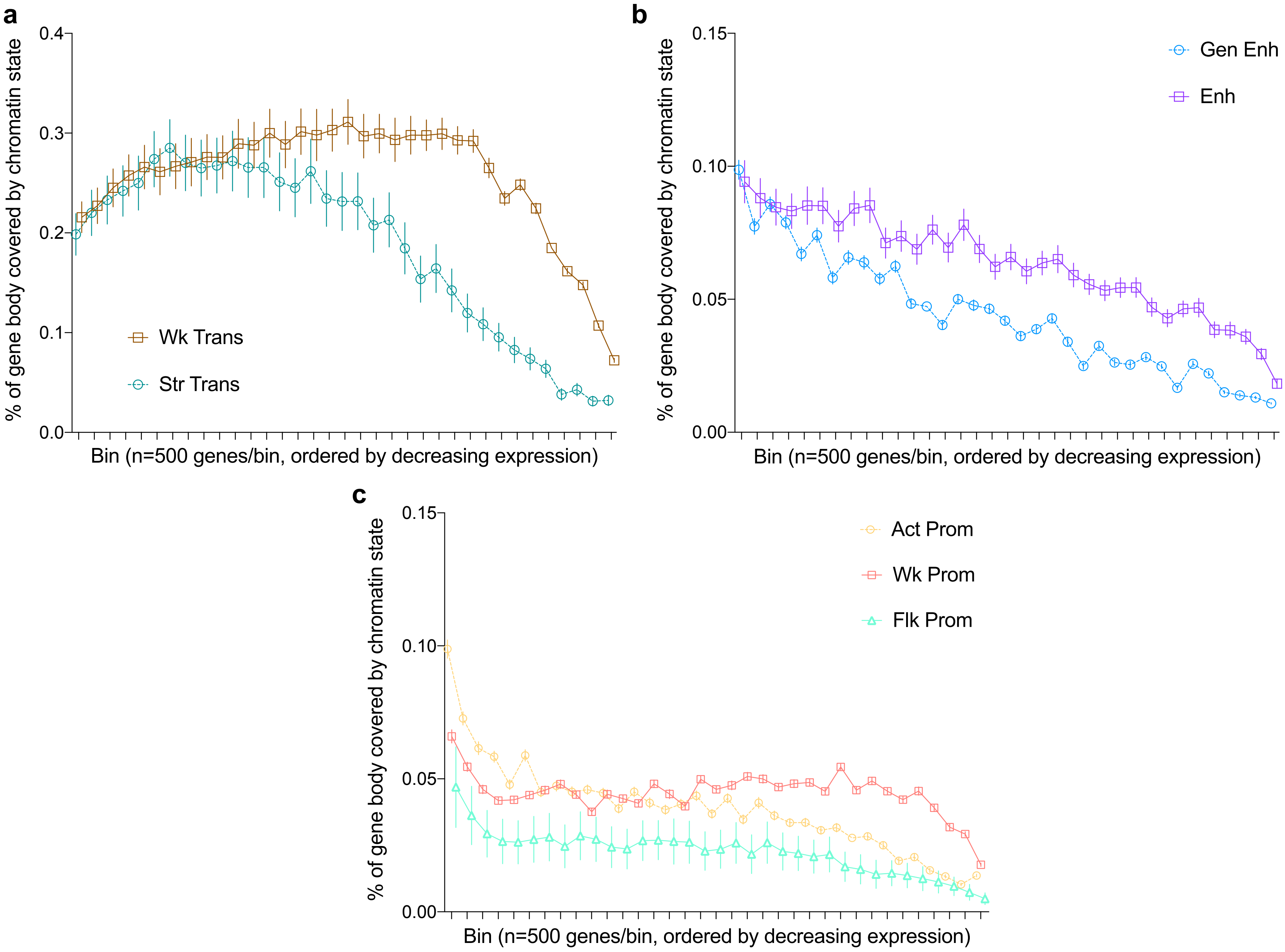
a
0.4
0.3
0.2
0.1
0.0
% of gene body covered by chromatin state
Wk Trans
Str Trans
Bin (n=500 genes/bin, ordered by decreasing expression)
b
0.15
0.10
0.05
0.00
% of gene body covered by chromatin state
Gen Enh
Enh
Bin (n=500 genes/bin, ordered by decreasing expression)
c
0.15
0.10
0.05
0.00
% of gene body covered by chromatin state
Act Prom
Wk Prom
Flk Prom
Bin (n=500 genes/bin, ordered by decreasing expression)

### FigureS5

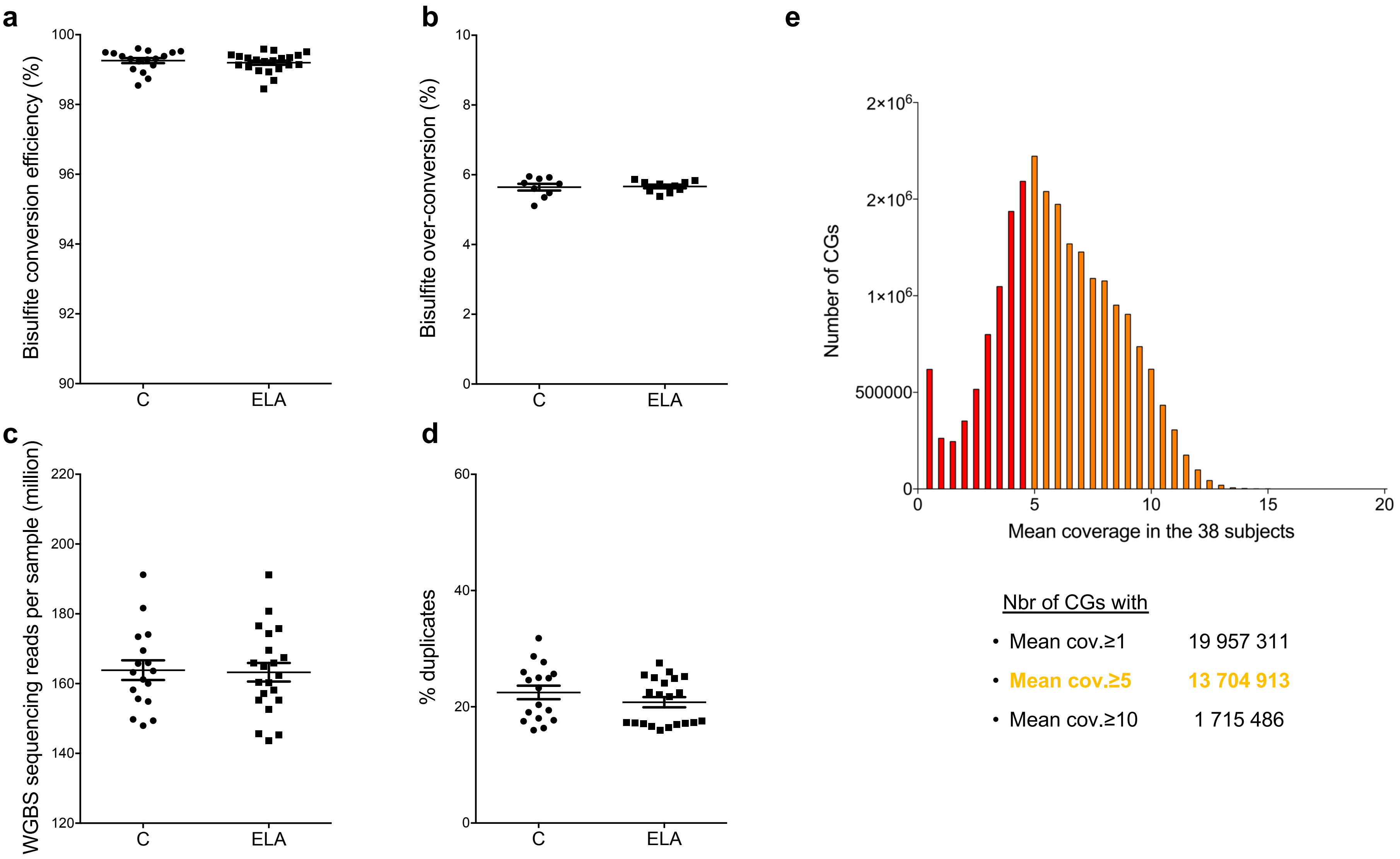

a
Bisulfite conversion efficiency (%)
100
98
96
94
92
90
C
ELA
b
Bisulfite over-conversion (%)
10
8
6
4
2
0
C
ELA
c
WGBS sequencing reads per sample (million)
220
200
180
160
140
120
C
ELA
d
% duplicates
60
40
20
0
C
ELA
e
Number of CGs
2×10⁶
2×10⁶
1×10⁶
500000
0
0
5
10
15
20
Mean coverage in the 38 subjects
Nbr of CGs with
• Mean cov.≥1 19 957 311
• Mean cov.≥5 13 704 913
• Mean cov.≥10 1 715 486

### FigureS6

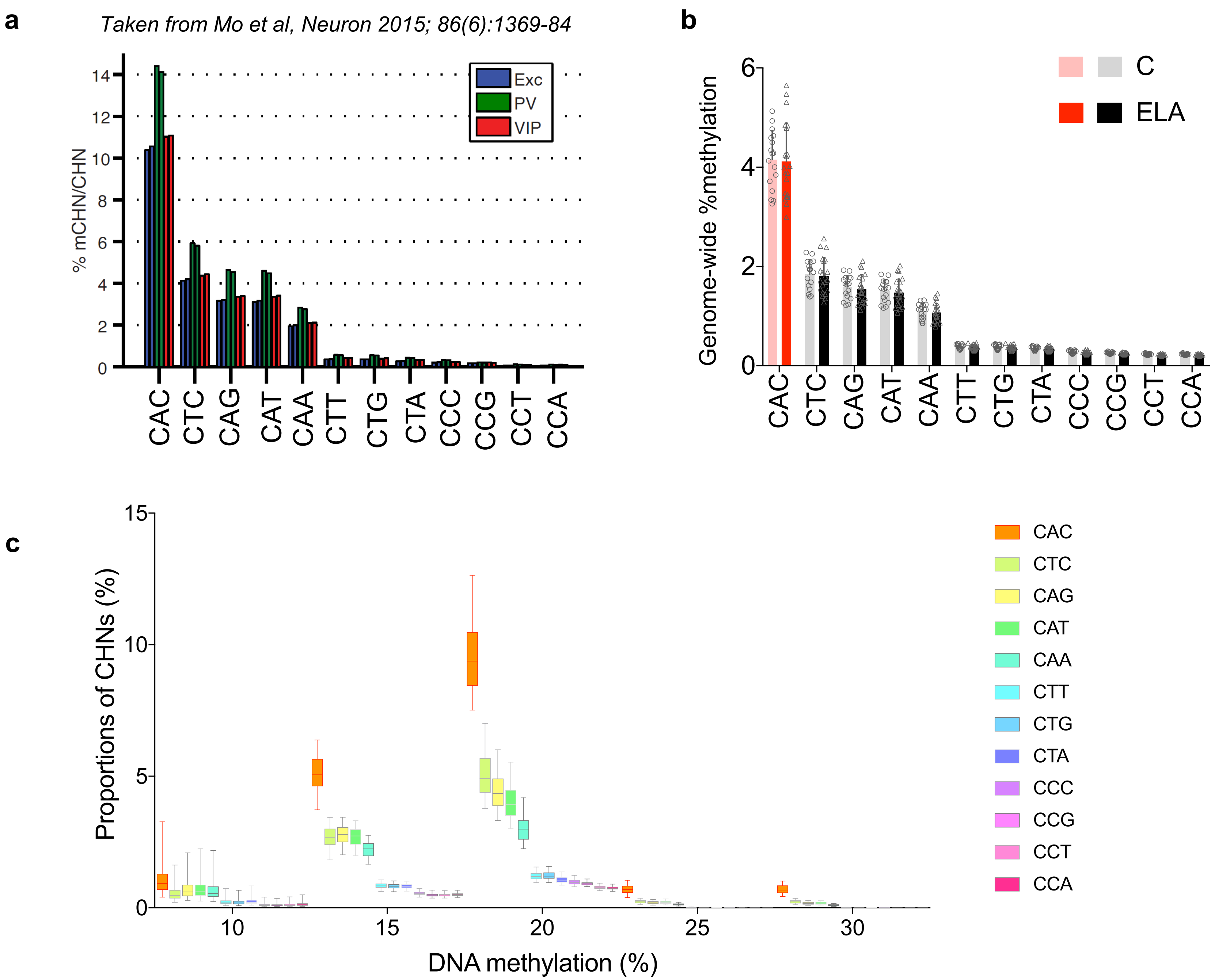
a
Taken from Mo et al, Neuron 2015; 86(6):1369-84
% mCHN/CHN
0
2
4
6
8
10
12
14
Exc
PV
VIP
CAC
CTC
CAG
CAT
CAA
CTT
CTG
CTA
CCC
CCG
CCT
CCA
b
Genome-wide %methylation
0
2
4
6
C
ELA
CAC
CTC
CAG
CAT
CAA
CTT
CTG
CTA
CCC
CCG
CCT
CCA
c
Proportions of CHNs (%)
0
5
10
15
DNA methylation (%)
10
15
20
25
30
CAC
CTC
CAG
CAT
CAA
CTT
CTG
CTA
CCC
CCG
CCT
CCA

FigureS7

**a**

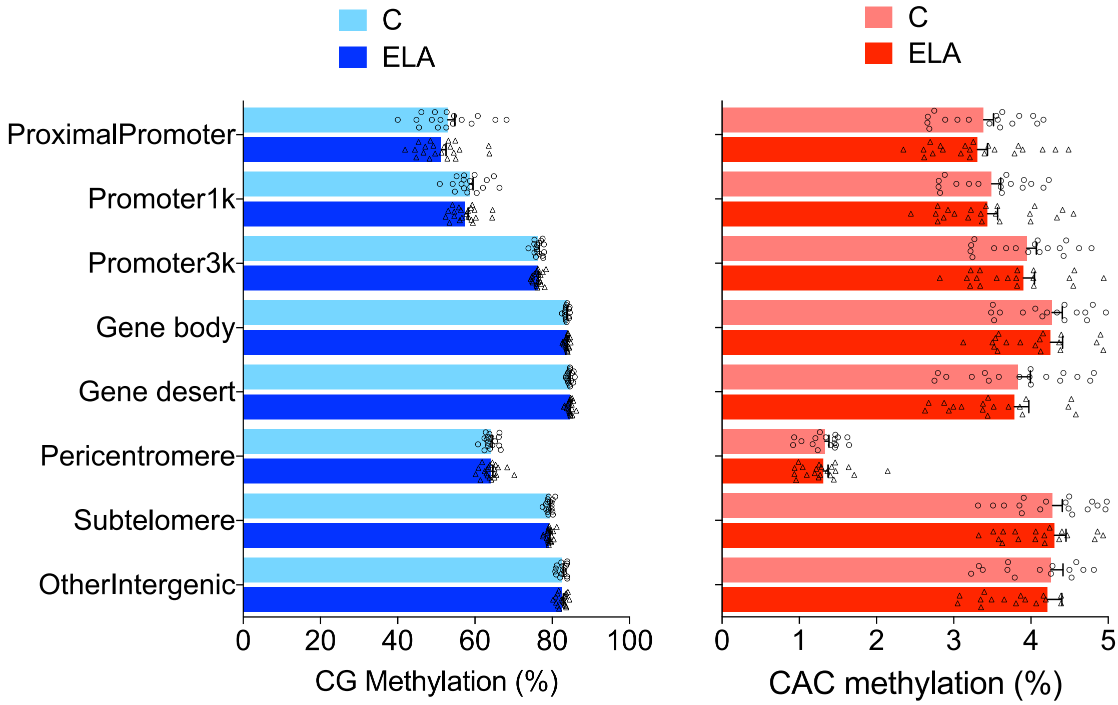
C
ELA
ProximalPromoter
Promoter1k
Promoter3k
Gene body
Gene desert
Pericentromere
Subtelomere
OtherIntergenic
0 20 40 60 80 100
CG Methylation (%)
C
ELA
0 1 2 3 4 5
CAC methylation (%)

**b**

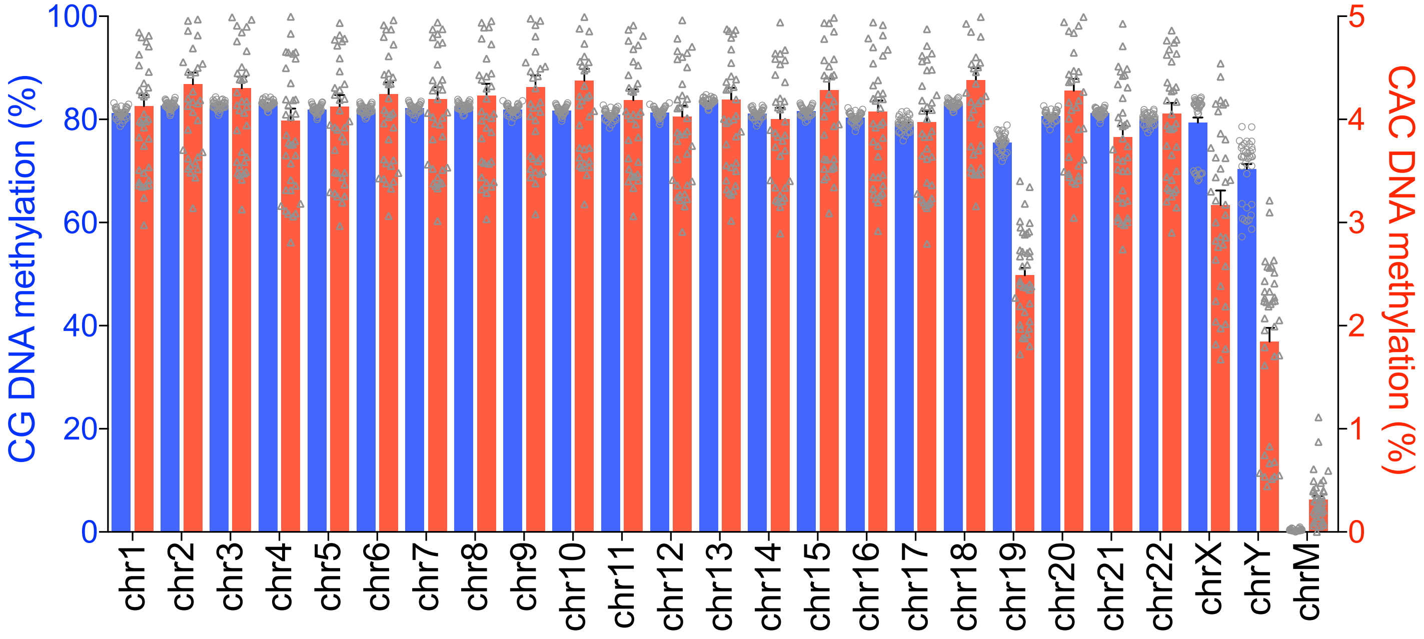
CG context
CAC context
CG DNA methylation (%)
0 20 40 60 80 100
CAC DNA methylation (%)
0 1 2 3 4 5
chr1 chr2 chr3 chr4 chr5 chr6 chr7 chr8 chr9 chr10 chr11 chr12 chr13 chr14 chr15 chr16 chr17 chr18 chr19 chr20 chr21 chr22 chrX chrY chrM

### FigureS8

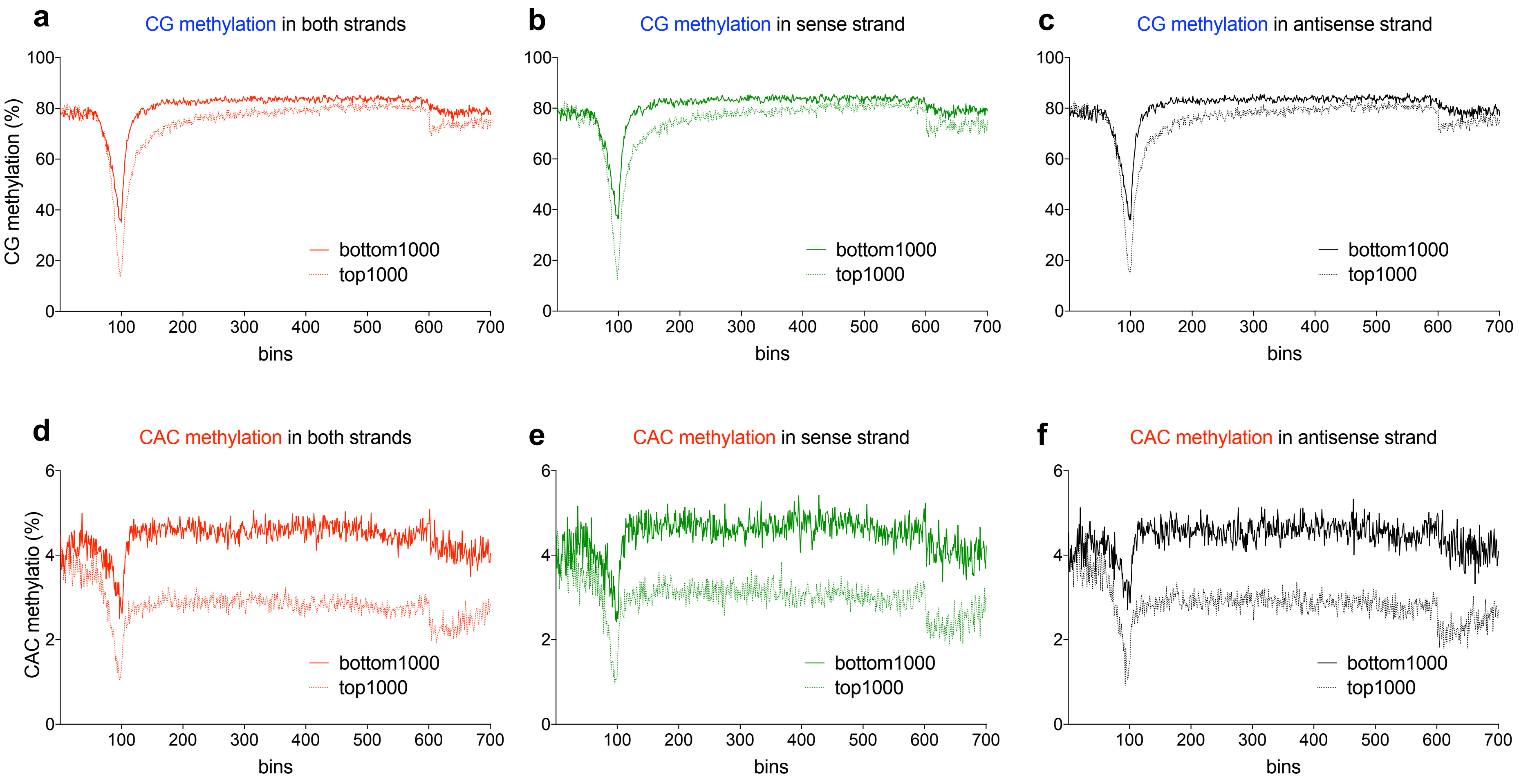
a
CG methylation in both strands
b
CG methylation in sense strand
c
CG methylation in antisense strand
CG methylation (%)
100
80
60
40
20
0
100 200 300 400 500 600 700
bins
bottom1000
top1000
d
CAC methylation in both strands
e
CAC methylation in sense strand
f
CAC methylation in antisense strand
CAC methylatio (%)
6
4
2
0

### FigureS9

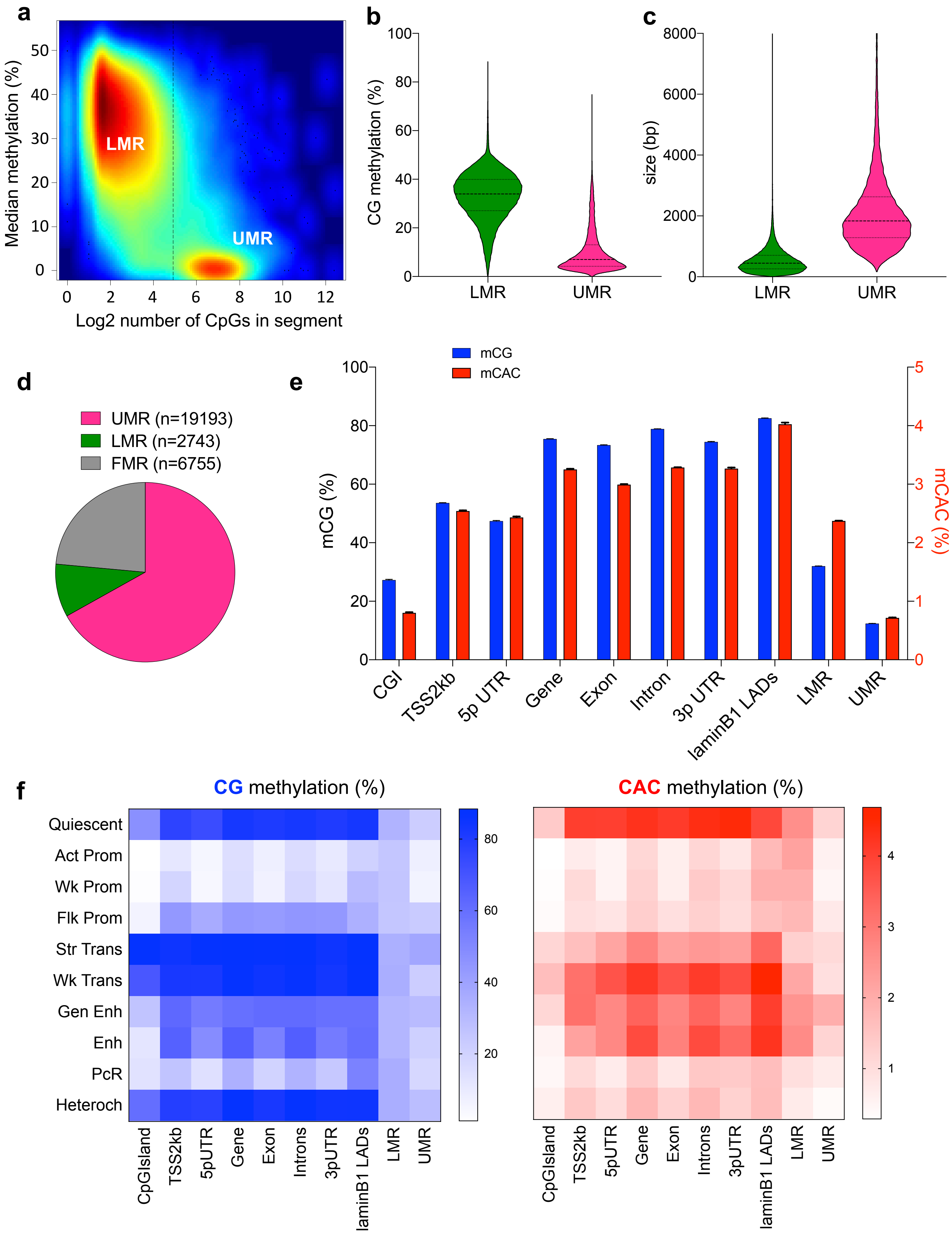
a
Median methylation (%)
Log2 number of CpGs in segment
LMR
UMR
b
CG methylation (%)
LMR
UMR
c
size (bp)
LMR
UMR
d
UMR (n=19193)
LMR (n=2743)
FMR (n=6755)
e
mCG
mCAC
mCG (%)
mCAC (%)
CGI
TSS2kb
5p UTR
Gene
Exon
Intron
3p UTR
laminB1 LADs
LMR
UMR
f
CG methylation (%)
CAC methylation (%)
Quiescent
Act Prom
Wk Prom
Flk Prom
Str Trans
Wk Trans
Gen Enh
Enh
PcR
Heteroch
CpGIsland
TSS2kb
5pUTR
Gene
Exon
Introns
3pUTR
laminB1 LADs
LMR
UMR

### FigureS10

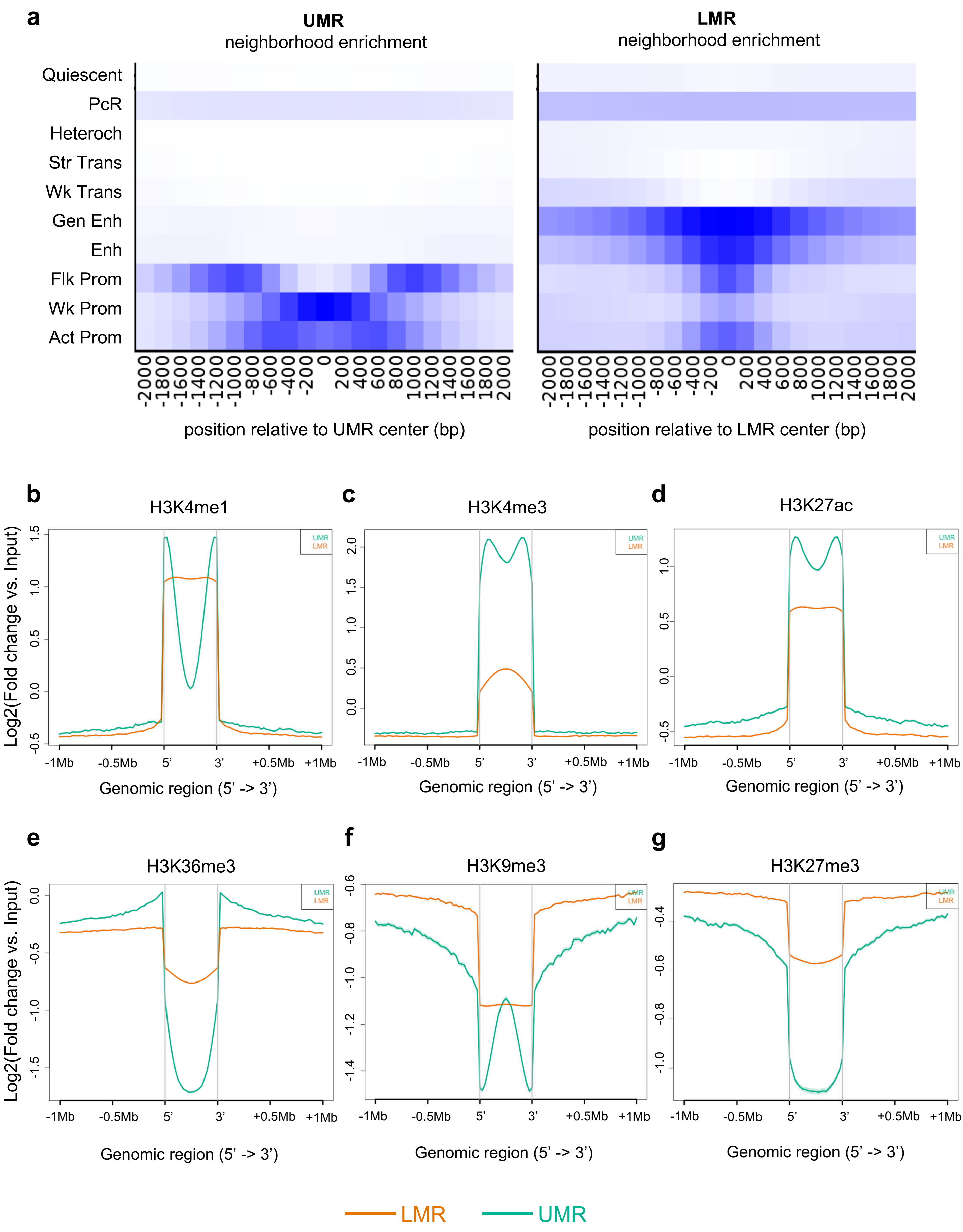
a
UMR
neighborhood enrichment
LMR
neighborhood enrichment
Quiescent
PcR
Heteroch
Str Trans
Wk Trans
Gen Enh
Enh
Flk Prom
Wk Prom
Act Prom
-2000 -1800 -1600 -1400 -1200 -1000 -800 -600 -400 -200 0 200 400 600 800 1000 1200 1400 1600 1800 2000
position relative to UMR center (bp)
position relative to LMR center (bp)
b
H3K4me1
c
H3K4me3
d
H3K27ac
e
H3K36me3
f
H3K9me3
g
H3K27me3
Log2(Fold change vs. Input)
-1Mb -0.5Mb 5' 3' +0.5Mb +1Mb
Genomic region (5' -> 3')
UMR
LMR

### FigureS11

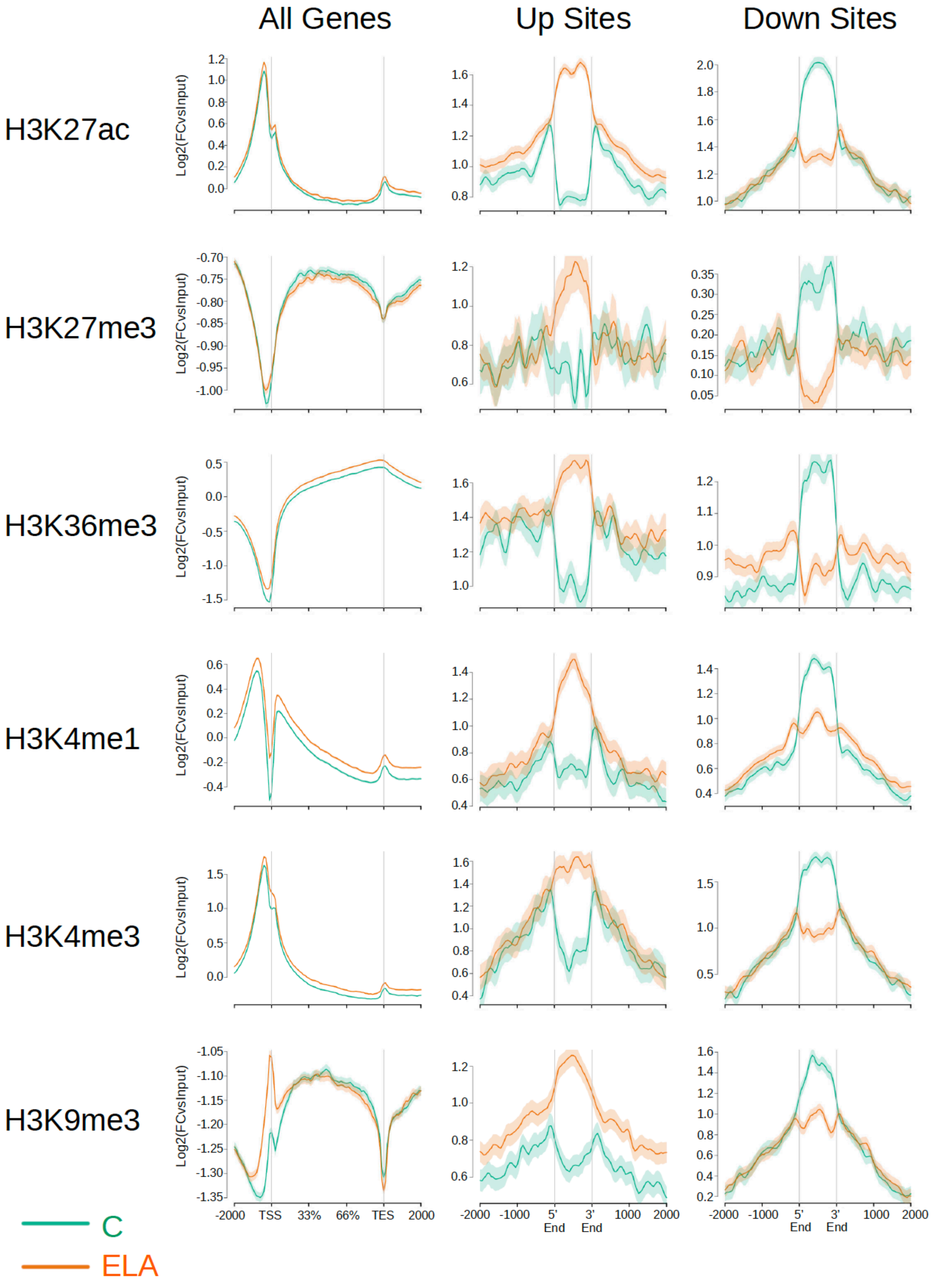

All Genes
Up Sites
Down Sites
H3K27ac
H3K27me3
H3K36me3
H3K4me1
H3K4me3
H3K9me3
Log2(FCvsInput)
-2000
TSS
33%
66%
TES
2000
-1000
5' End
3' End
1000
C
ELA

### FigureS12

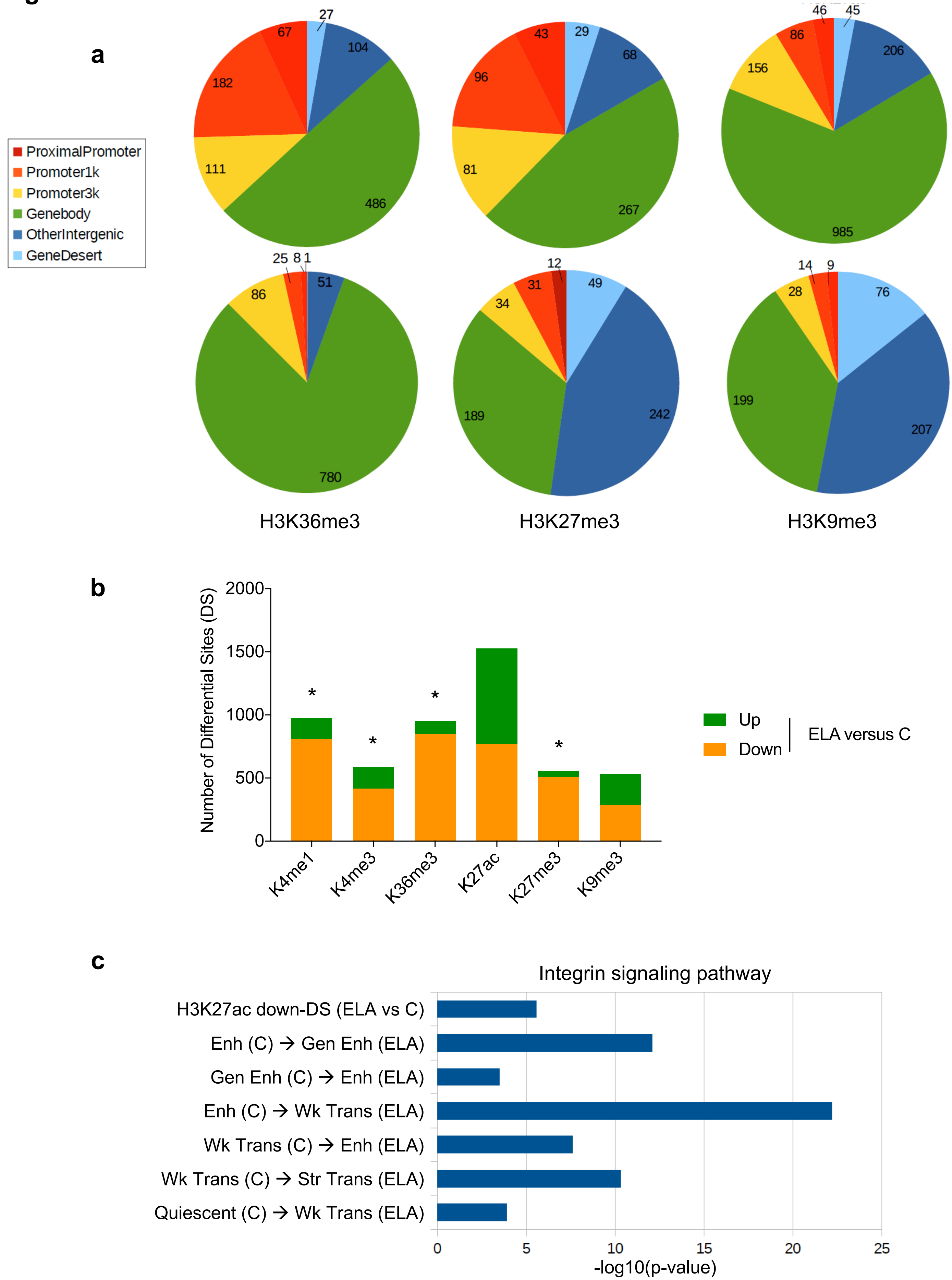
a
H3K4me1
H3K4me3
H3K27ac
H3K36me3
H3K27me3
H3K9me3
ProximalPromoter
Promoter1k
Promoter3k
Genebody
OtherIntergenic
GeneDesert
27
104
486
111
182
67
29
68
267
81
96
43
46
45
206
985
156
86
25 8 1
51
780
86
12
49
242
189
34
31
14 9
76
207
199
28
b
Number of Differential Sites (DS)
2000
1500
1000
500
0
K4me1
K4me3
K36me3
K27ac
K27me3
K9me3
*
Up
Down
ELA versus C
c
Integrin signaling pathway
H3K27ac down-DS (ELA vs C)
Enh (C) → Gen Enh (ELA)
Gen Enh (C) → Enh (ELA)
Enh (C) → Wk Trans (ELA)
Wk Trans (C) → Enh (ELA)
Wk Trans (C) → Str Trans (ELA)
Quiescent (C) → Wk Trans (ELA)
0
5
10
15
20
25
-log10(p-value)

**FigureS13**

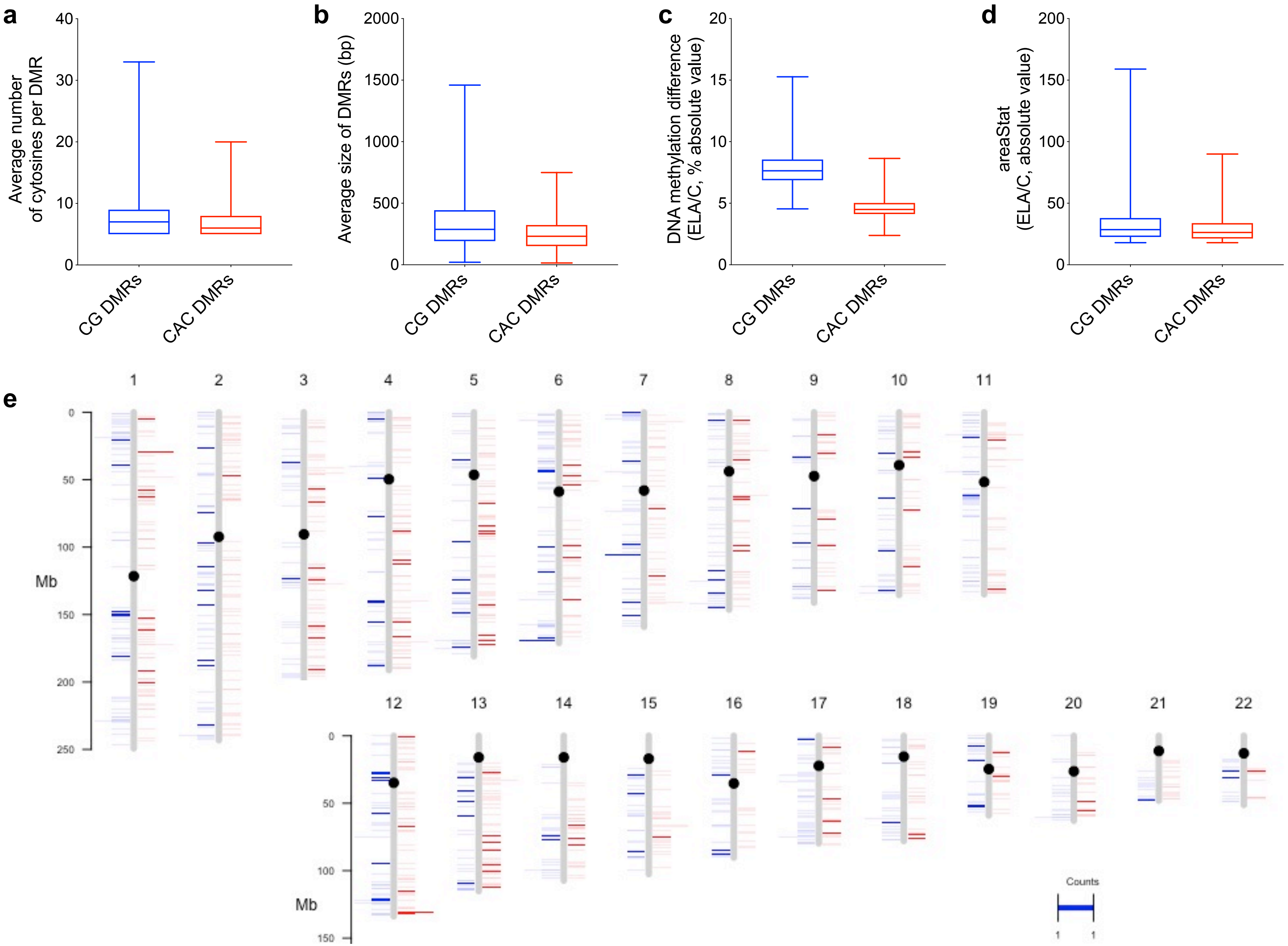
a
Average number of cytosines per DMR
0
10
20
30
40
CG DMRs
CAC DMRs
b
Average size of DMRs (bp)
0
500
1000
1500
2000
CG DMRs
CAC DMRs
c
DNA methylation difference (ELA/C, % absolute value)
0
5
10
15
20
CG DMRs
CAC DMRs
d
areaStat (ELA/C, absolute value)
0
50
100
150
200
CG DMRs
CAC DMRs
e
1 2 3 4 5 6 7 8 9 10 11
0
50
100
150
200
250
Mb
12 13 14 15 16 17 18 19 20 21 22
0
50
100
150
Mb
Counts
1
1

### FigureS14

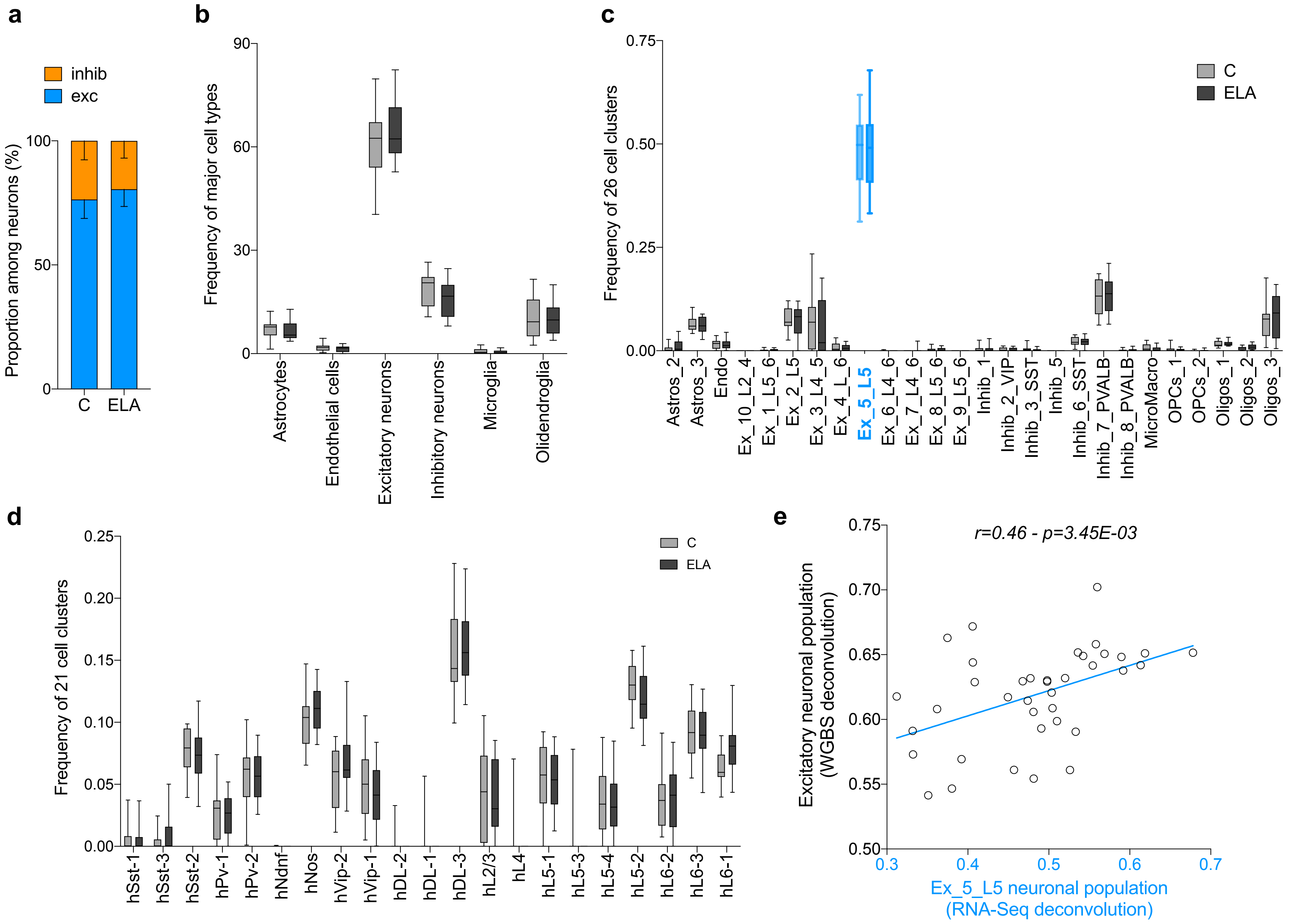

a
inhib
exc
Proportion among neurons (%)
100
50
0
C
ELA
b
Frequency of major cell types
90
60
30
0
Astrocytes
Endothelial cells
Excitatory neurons
Inhibitory neurons
Microglia
Olidendroglia
c
Frequency of 26 cell clusters
0.75
0.50
0.25
0.00
C
ELA
Astros_2
Astros_3
Endo
Ex_10_L2_4
Ex_1_L5_6
Ex_2_L5
Ex_3_L4_5
Ex_4_L_6
Ex_5_L5
Ex_6_L4_6
Ex_7_L4_6
Ex_8_L5_6
Ex_9_L5_6
Inhib_1
Inhib_2_VIP
Inhib_3_SST
Inhib_5
Inhib_6_SST
Inhib_7_PVALB
Inhib_8_PVALB
MicroMacro
OPCs_1
OPCs_2
Oligos_1
Oligos_2
Oligos_3
d
Frequency of 21 cell clusters
0.25
0.20
0.15
0.10
0.05
0.00
C
ELA
hSst-1
hSst-3
hSst-2
hPv-1
hPv-2
hNdnf
hNos
hVip-2
hVip-1
hDL-2
hDL-1
hDL-3
hL2/3
hL4
hL5-1
hL5-3
hL5-4
hL5-2
hL6-2
hL6-3
hL6-1
e
r=0.46 - p=3.45E-03
Excitatory neuronal population (WGBS deconvolution)
0.75
0.70
0.65
0.60
0.55
0.50
0.3
0.4
0.5
0.6
0.7
Ex_5_L5 neuronal population (RNA-Seq deconvolution)

### FigureS15

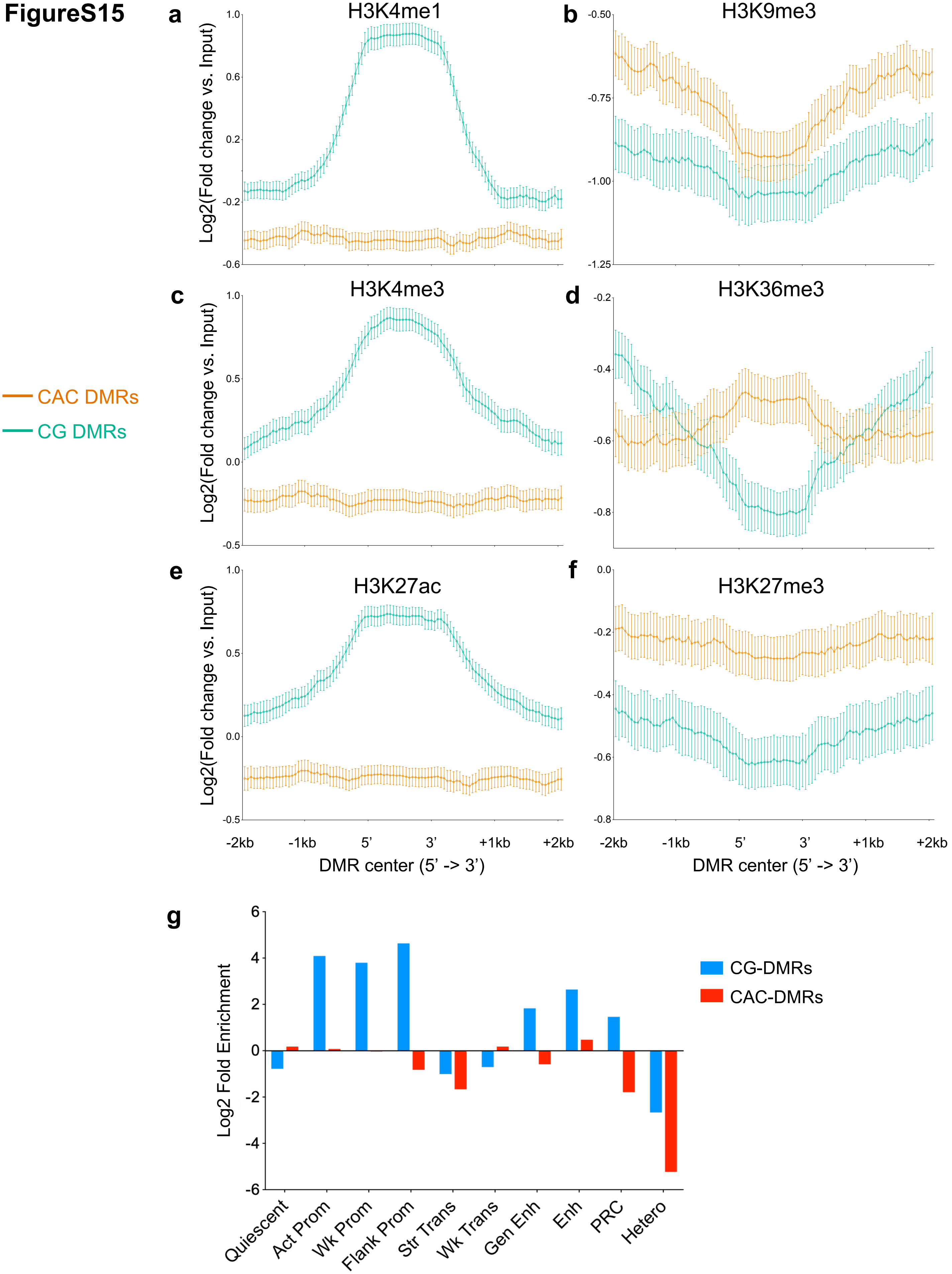

a
H3K4me1
b
H3K9me3
c
H3K4me3
d
H3K36me3
e
H3K27ac
f
H3K27me3
Log2(Fold change vs. Input)
-2kb
-1kb
5'
3'
+1kb
+2kb
DMR center (5' -> 3')
CAC DMRs
CG DMRs
g
Log2 Fold Enrichment
CG-DMRs
CAC-DMRs
Quiescent
Act Prom
Wk Prom
Flank Prom
Str Trans
Wk Trans
Gen Enh
Enh
PRC
Hetero

**FigureS16**

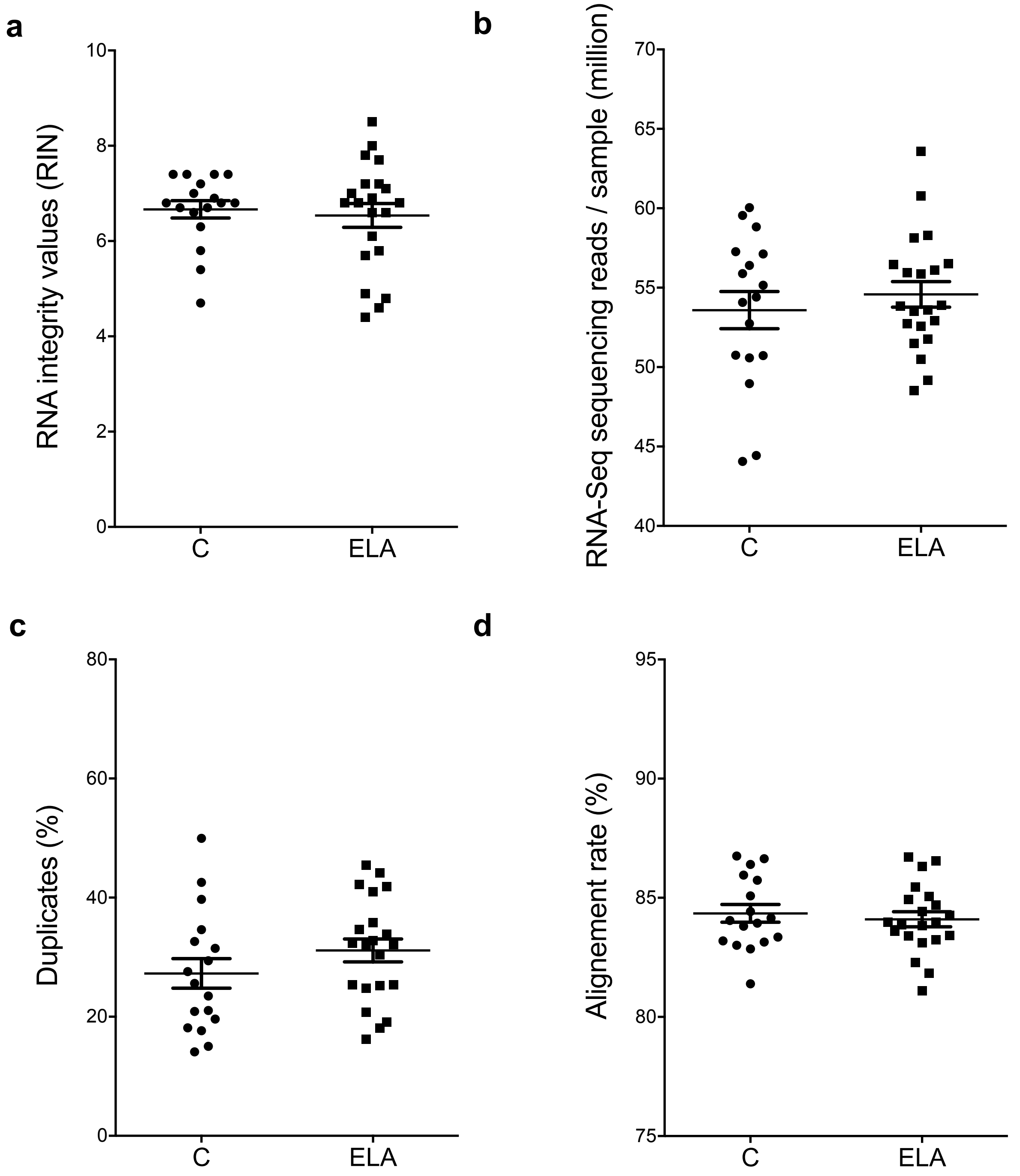
a
b
c
d
RNA integrity values (RIN)
RNA-Seq sequencing reads / sample (million)
Duplicates (%)
Alignement rate (%)
0
2
4
6
8
10
40
45
50
55
60
65
70
0
20
40
60
80
75
80
85
90
95
C
ELA

**FigureS17**

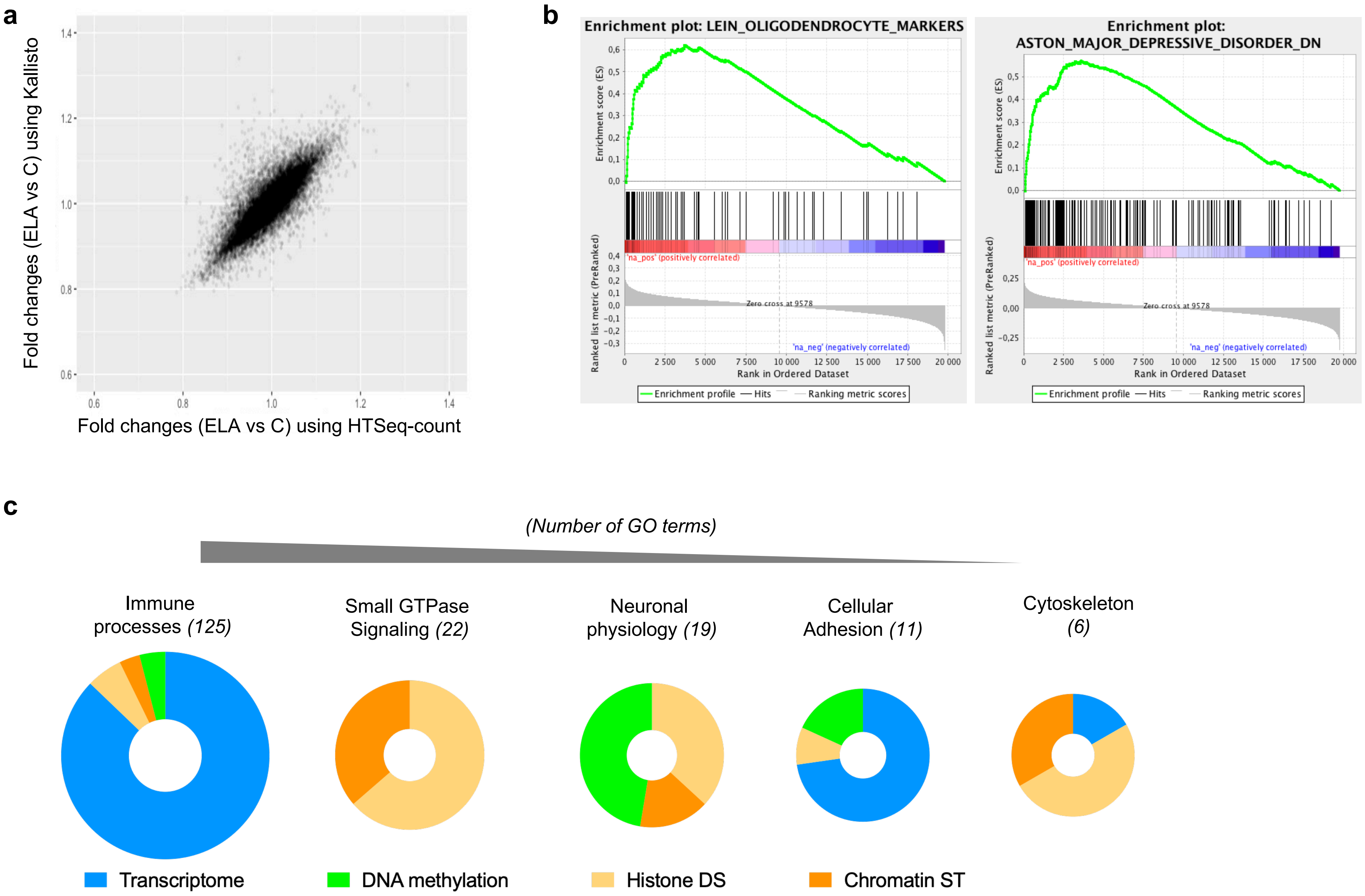
a
Fold changes (ELA vs C) using Kallisto
Fold changes (ELA vs C) using HTSeq-count
b
Enrichment plot: LEIN_OLIGODENDROCYTE_MARKERS
Enrichment plot: ASTON_MAJOR_DEPRESSIVE_DISORDER_DN
Enrichment score (ES)
Ranked list metric (PreRanked)
'na_pos' (positively correlated)
Zero cross at 9578
'na_neg' (negatively correlated)
Rank in Ordered Dataset
Enrichment profile
Hits
Ranking metric scores
c
(Number of GO terms)
Immune processes (125)
Small GTPase Signaling (22)
Neuronal physiology (19)
Cellular Adhesion (11)
Cytoskeleton (6)
Transcriptome
DNA methylation
Histone DS
Chromatin ST
